# Control of cell fate specification and patterning by an ancestral microRNA

**DOI:** 10.1101/2023.09.09.556951

**Authors:** Adolfo Aguilar-Cruz, Eduardo Flores-Sandoval, Ximena Gutiérrez-Ramos, Omar Oltehua- Lopez, Ana E. Dorantes-Acosta, Joshua T. Trujillo, Hirotaka Kato, Kimitsune Ishizaki, Rebecca A. Mosher, Liam Dolan, Daniel Grimanelli, Jim Haseloff, John L. Bowman, Mario A. Arteaga-Vazquez

## Abstract

The formation of an organized body requires the establishment and maintenance of cells with structural and functional distinctive characteristics. A central question in developmental biology is how changes in the regulation of genes drive cell specification and patterning^1^. microRNAs (miRNAs) are small non-coding RNAs that regulate development through mRNA cleavage and/or translational repression^2^. In plants, miRNAs regulate key aspects including growth, development, stem cell maintenance, vegetative phase change, leaf morphogenesis, floral organ formation and flowering time^3^. Biogenesis of plant miRNAs depends on the activity of DICER-LIKE 1 (DCL1), an RNase type III endonuclease that processes double stranded RNA to give rise to mature miRNAs ^4^. The genomes of today’s flora contain at least one bona fide copy of *DCL1* ^5,6^. Using *Marchantia polymorpha* -a model bryophyte that allows comparative approaches to infer characteristics of the ancestral land plant-, we demonstrate that Mp*DCL1a* is required for the biogenesis of miRNAs and uncovered a central role for miR166/Homeodomain Zipper Class III-regulated auxin synthesis in the specification of cell identity, patterning, meristem function, laminar expansion and the development of the body in the last common ancestor of extant land plants.

## INTRODUCTION

MicroRNAs (miRNAs) are small non-coding RNA molecules ∼21 nucleotides in length that regulate plant development and responses to internal and external environmental cues^3^. MiRNAs regulate gene expression post-transcriptionally by a mechanism that depends on the sequence complementarity between the miRNA and its target mRNA^2^. In plants, the biogenesis of miRNAs requires the activity of specialized RNase type III enzymes known as DICER-LIKE (DCL) proteins ^7^. All members of the canonical DCL family contain six evolutionarily conserved functional domains: N-terminal DExD/H-box (pfam: PF00270), Helicase (PF00271), DUF283 (PF03368), PAZ (PF02170), two RNase III domains (PF00636), and a dsRNA-binding domain (dsRBD) (PF14709) ^8^. DCL1 is the major enzyme involved in the biogenesis of 20-21 nt miRNAs ^4^. DCL1 is an essential gene in Arabidopsis (*dcl1* null alleles are embryo lethal)^9^, indicating that no other member of the DCL family can complement its function. The presence of miRNAs and members of the DCL1 family occurs across the Viridiplantae clade including extant basal land plant lineages^6^ (*i.e.,* bryophytes and tracheophytes).

*Marchantia polymorpha* is a liverwort that belongs to the group of non-vascular plants -the bryophytes-that diverged from the ancestor of vascular plants and angiosperms over 470 million years ago^10,11^. The basic body plan of the dominant gametophytic phase of *M. polymorpha* is a dorsiventral thallus that grows as a flat sheet that follows a characteristic bifurcation pattern that results from the periodic duplication of meristems^12^. Gemma cups producing asexual vegetative propagules (gemmae) are formed on the dorsal side^13^. Gemmae are multicellular bilobulated propagules that contain two meristematic growing points also known as apical notches that through symmetrical growth and divisions develop into a thallus^14^. Gemmae development initiates when an epidermal cell of the floor of a gemma cup undergoes an asymmetrical division to produce a gemma initial that will further divide to produce a proximal and a distal cell. The distal cell divides to produce the gemma body and the proximal cell enlarges without further divisions to form the stalk cell, that will connect the gemma body to the floor of the gemma cup^14–17^ (Figure 1a). Hormones auxin and cytokinin are involved in the formation and development of gemma cups and gemmae^18,19^. In this work, we show that Mp*DCL1a* is a pleiotropic gene that controls diverse developmental programs including cell fate specification, patterning, cell division, growth and phase transition. We found that Mp*DCL1a* is the major contributor for the biogenesis of miRNAs and mutations in this locus cause accumulation of miRNA precursors and miRNA targets. In addition, Mp*dcl1a* loss-of-function mutants resemble the phenotype of miRNA loss-of-function mutants. Finally, we demonstrate that the miR166-C3HDZ module regulates patterning cooperatively with auxin, cell identity, meristem function, laminar expansion and controls asexual reproduction through the specification of gemma cell fate.

**Figure 1.**
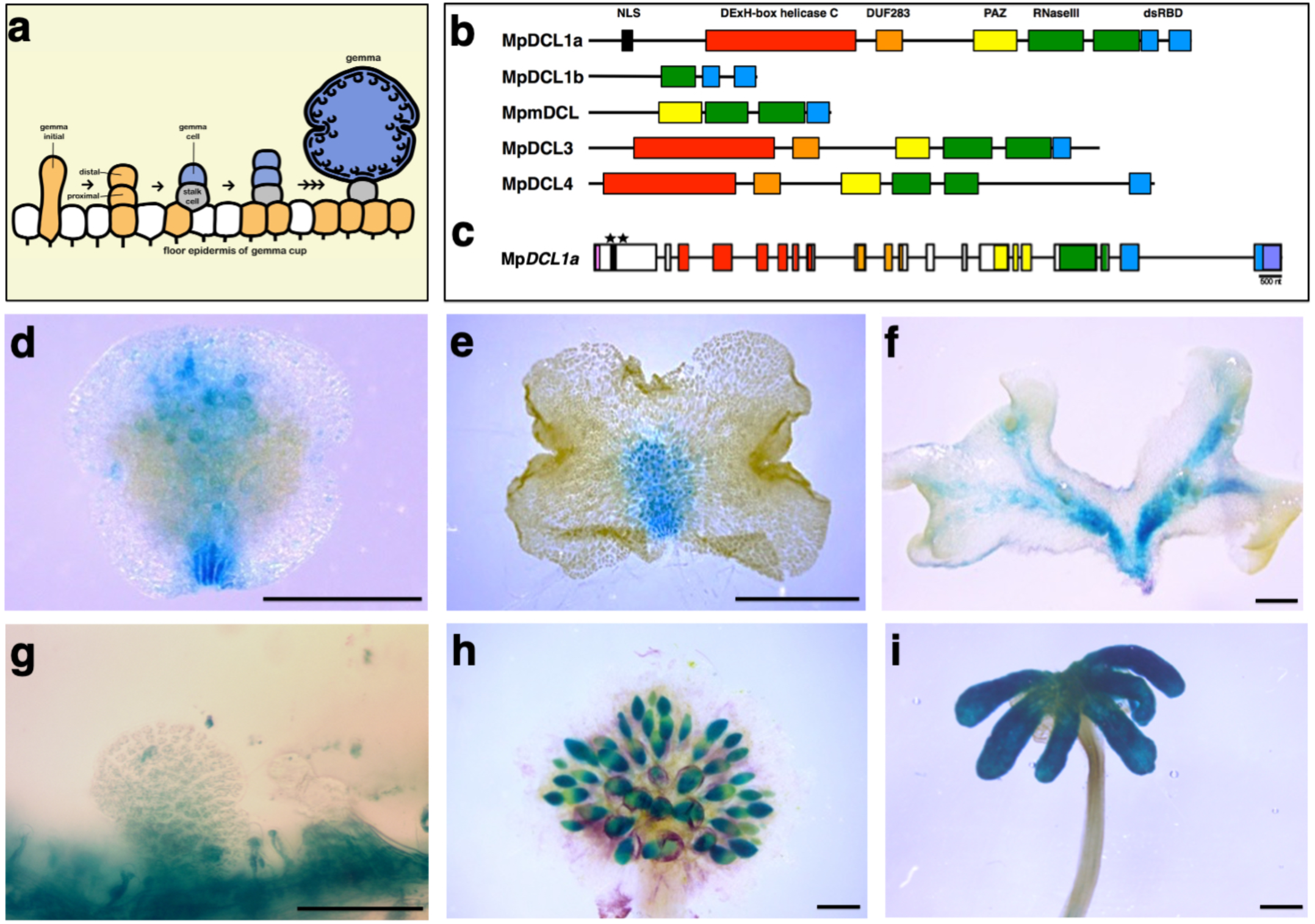
Formation of gemma and composition of the DCL family in *Marchantia polymorpha.* a) Diagram depicting the gemmae development from the floor epidermis of gemma cups. b) Phylogenetic classification of the DCL family in *Marchantia polymorpha*. The evolutionarily conserved functional domains are depicted as colored boxes: DExH-box helicase C (red), DUF283 (orange), PAZ (yellow), RNaseIII (green) and dsRBD (blue). A nuclear localization signal (NLS) present in MpDCL1a is shown as a black box. c) Genomic structure of Mp*DCL1a*, introns are shown as black lines and exons as boxes. Colored boxes depict particular exons encoding functional domains and follow the same coloring pattern as in panel b. The genomic locations of the two gRNAs targeting MpDCL1a are indicated with asterisks. d-i) Expression pattern of Mp*DCL1a* at different developmental stages. GUS staining is observed in d) the central zone of gemma and the cells formerly attached to the maternal tissue, e) the central zone of 2-days-old gemmalings, f) the floor of gemma cups, g) the midrib of 21-days-old thalli, h) antheridiophores, i) archegoniophores. Scale bars: d and f = 0.5 mm. e= 0.1 mm. g-i = 1 mm.

## RESULTS

### The *Marchantia polymorpha* genome encodes five *DICER-LIKE* genes

The *M. polymorpha* genome was previously shown to encode four members of the DCL family: MpDCL1a (Mp7g12090.1), MpDCL1b (Mp7g16380.1), MpDCL3 (Mp1g02840.1) and MpDCL4 (Mp7g11720.1) with MpDCL1b being atypical since it contained an incomplete set of the signature domains. We reexamined the annotation of the *DCL* family in *M. polymorpha* and identified a *M. polymorpha MINIMAL DICER-LIKE* (Mp*mDCL*) (Mp6g09830.1) gene (Figure 1b). All evolutionarily conserved signature domains of the DCL family are present in MpDCL1a, MpDCL3 and MpDCL4 but some are absent in the non-canonical MpDCL1b and MpmDCL (Figure 1b and S1). These results indicate that in addition to the three previously reported clades (DCL1, DCL3 and DCL4)^10^, the *M. polymorpha* genome encodes a fourth clade containing a non-canonical MpmDCL (Figure 1b and S1).

### Mp*DCL1a* is expressed across the *M. polymorpha* life cycle

The Mp*DCL1a* gene (Mp7g12090.1) has a length of 15,352 nt (Figure 1c) and encodes a 2,057 amino acids protein (Figure 1b). Mp*DCL1*a has 20 exons and 19 introns, it shows a 67.7% identity with *Arabidopsis thaliana* DCL1 and encodes a protein with all the conserved functional domains of the DCL family^10^ (Figure 1b). The expression pattern of Mp*DCL1a* that we obtained by RT-PCR (*i.e.* gemmae, mature thalli and gametangiophores) (Figure S2a) is consistent with the transcriptomic profile reported for Mp*DCL1a* in Marpol Base^20^ (https://marchantia.info/mbex/). We generated transgenic *M. polymorpha* lines harboring a transcriptional fusion of a 5 kb genomic fragment upstream of the ATG codon (including the 5’UTR region) of Mp*DCL1a*, driving the expression of the beta-glucuronidase (*GUS*) reporter gene. We observed GUS staining predominantly in the central zone of the gemma encompassing the rhizoid initiation zone and the cells formerly attaching the gemma to the stalk cell present in the maternal tissue of the gemma cup (Figure 1d) and papillae cells (Figure S2b). In young thalli, GUS staining was observed in the central region (Figure 1e). In mature thalli, GUS staining was localized across the midrib and to the surroundings of the apical notch (but not in the apical notch) (Figure 1f). In gemma cups, *GUS* expression was observed in the basal floor of the cup and developing gemmae (Figure 1g). Strong GUS staining was observed in antheridiophores and antheridia (Figure 1h), archegoniophores and digitate rays (Figure 1i), venter, collar cells and the egg cell (Figure S2c). Taken together, this indicates that Mp*DCL1a* is expressed across the life cycle of *M. polymorpha* including thalli, gemminiferous tissues, gemmae, gametangiophores and gametes.

### Generation of Mp*DCL1a* partial loss-of-function mutants

We selected two guide RNAs (gRNAs) based on the presence of a methionine codon in frame with the CDS of Mp*DCL1a* located 356 bp apart from each other, targeting the sense strand of the first exon of Mp*DCL1a* (Figure 1c). Despite intensive efforts, we have not been able to isolate Mp*dcl1a* complete loss-of-function alleles which strongly suggest that Mp*DCL1a* is also an essential gene, as observed in other plants^9^. We obtained more than 100 independent events that were subsequently screened for mutations within the Mp*DCL1a* locus by PCR amplification and DNA sequencing. We detected mutations ranging from single nucleotide insertions/deletions to deletions of up to 511 nucleotides (Figure 2a). All recovered Mp*DCL1a* mutant alleles disrupted a genomic region encoding a nuclear localization signal (NLS) but did not affect either its transcription nor the evolutionarily conserved domains (Figure 1c and 2a).

**Figure 2.**
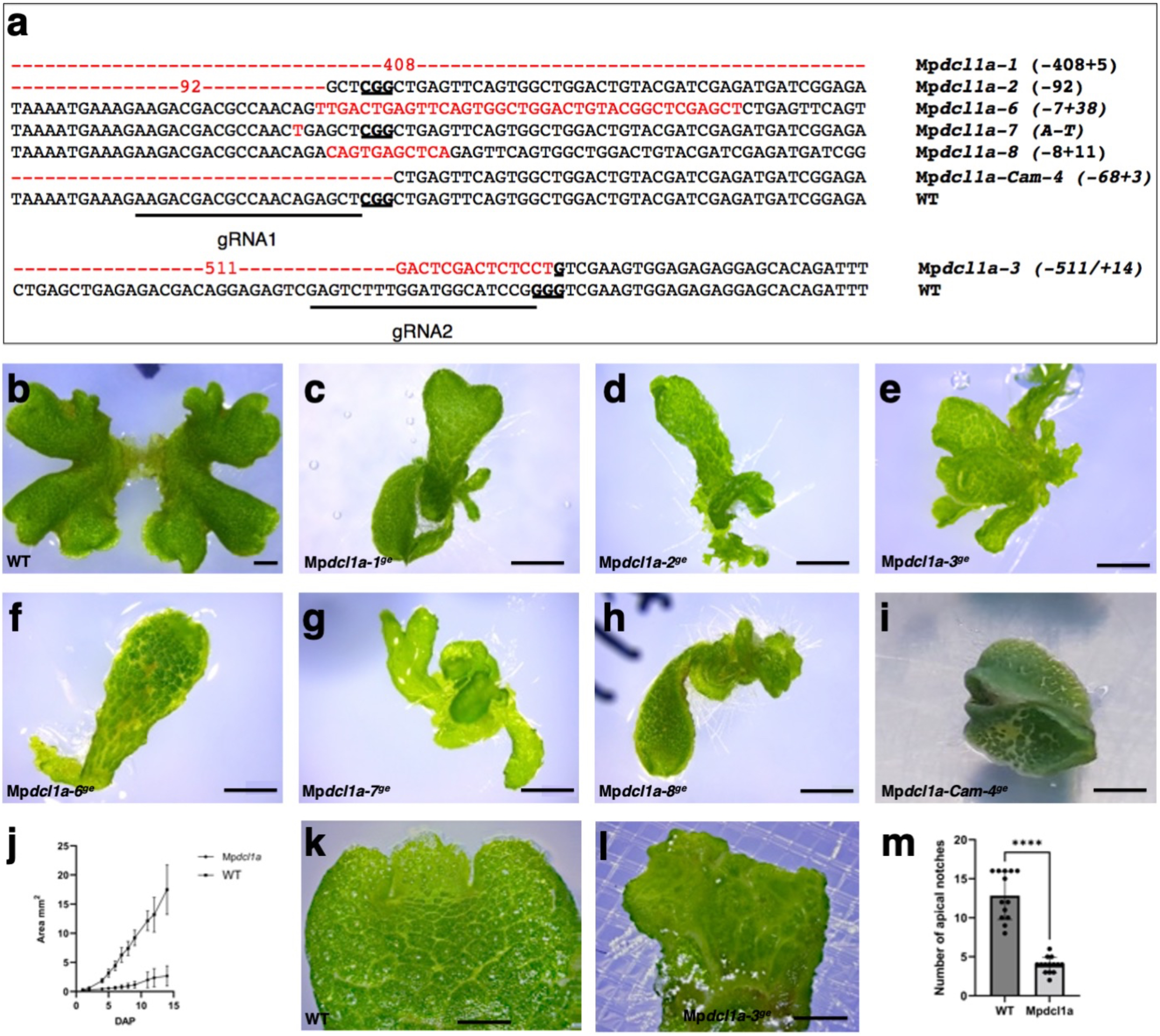
The partial loss-of-function of Mp*DCL1a* affects growth and development. a) Summary of genomic edits generated in Mp*DCL1a* using gRNA1 and 2 (underlined sequences). Insertions and deletions (indels) for each mutant allele are indicated in parenthesis and depicted in red font. PAM sequences are shown as bold underlined letters. b-i) Phenotype of 21-day-old wild type (WT) and Mp*dcl1a* loss-of-function plants. b) WT, c) Mp*dcl1a-1^ge^*, d) Mp*dcl1a-2^ge^*, e) Mp*dcl1a-3^ge^*, f) Mp*dcl1a-6^ge^*, g) Mp*dcl1a-7^ge^*, h) Mp*dcl1a-8^ge^* and i) Mp*dcl1a-Cam-4^ge^*. j) Growth rate of WT and Mp*dcl1a-3^ge^* plants. k) Branching phenotype in 13-day-old WT. l) Branching phenotype in 13-day-old Mp*dcl1a-3^ge^* plants. m) Quantification of the number of apical notches in WT and Mp*dcl1a-3^ge^*plants based on the expression pattern of the apical meristem marker _pro_Mp*YUC2:GUS* (Figure S3). Scale bars: b-i = 1 mm. j and k = 0.5 mm.

### Mp*DCL1a* is a pleiotropic gene that controls growth and development

Size and growth of Mp*dcl1a* mutants are defective, their thalli are elongated and narrowed laterally and therefore tongue-shaped (Figure 2c-i). Fourteen day-old Mp*dcl1a* mutants were on average nine times shorter than wild-type plants (Figure 2j). To determine the pattern of development of Mp*dcl1a* mutants, we scored the timing of the periodic duplication of meristems (defined as the notch plastochron) as previously reported^12^. To quantify the number of apical notches we introduced the apical notch marker *pro*Mp*YUC2:GUS* ^21^ into the Mp*dcl1a* mutant background. Nine days after cultivation, wild-type plants are in the second plastochron, they exhibit on average four apical notches and the second division of the apical notches has just begun (Figure 2b, 2k, 2m and S3). By contrast, Mp*dcl1a* plants are in the first plastochron as apical notches just started dividing (Figure 2l-m and S3). Mp*dcl1a* plants develop fewer apical notches than wild type as the cells proliferate and the apical notch expands laterally but division and formation of the central lobe is delayed relative to wild type (Figure S3). These observations indicate that Mp*DCL1a* is required for thallus growth and development including the patterning and timing of division of the meristematic apical notch.

### Mp*DCL1a* regulates asexual reproduction by repressing the gemmae cell fate

Gemmae from hypomorphic Mp*dcl1a* mutant alleles with mutations in the first exon that eliminate the NLS, form supernumerary concave meristem-like structures and supernumerary papillae (Figure 3d-l). However, the most striking phenotype we observed was the formation of gemmae directly from the laminar surface of the gemma epidermis (Figure 3d-l). In dormant wild-type gemmae, dorsal and ventral epidermis are flat, and papillae are the only structures that expand out of the epidermal plane (Figure 3a-c). Mp*dcl1a* mutant gemmae develop ectopic gemmae directly from both the dorsal and ventral epidermis (Figure 3d-l). The formation of ectopic gemmae in Mp*dcl1a* occurs early in development because they can be observed in gemmae present within the gemma cup (Figure S4). Mp*dcl1a* mutant gemmae exhibit ectopic gemmae at different developmental stages ranging from cells resembling gemma initials to mature gemmae that also developed supernumerary papillae (Figure S4). This is reminiscent of the developmental series normally observed at the floor of gemma cups during the formation of gemmae. From a group of Mp*dcl1a* hypomorphs, we performed a detailed phenotypic and molecular characterization of the allele Mp*dcl1a-3^ge^*as the rest of the alleles either stopped producing gemmae or died shortly after forming reproductive structures without exposure to far-red light. These findings indicate that Mp*DCL1a* is not only required for the formation of gemmae but also to repress gemma cell fate and the formation of supernumerary papillae on the epidermis of the gemma.

**Figure 3.**
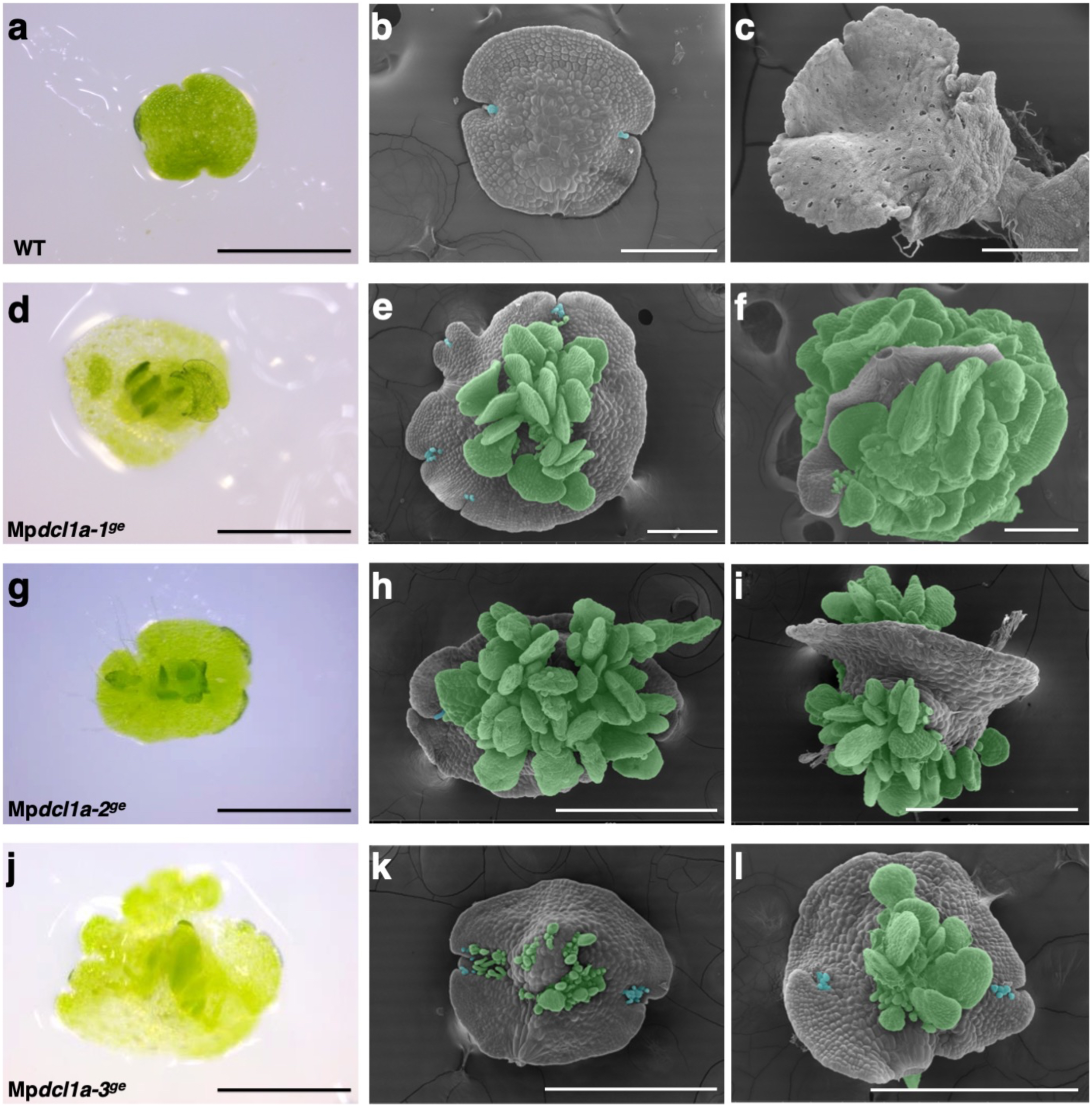
Patterning and cell specification defects in Mp*dcl1a* hypomorphic alleles. Bright field microscopy (BFM) and scanning electron microscopy (SEM) images of wild type (WT) and Mp*dcl1a* mutant alleles. a-c) BFM image of a WT gemma, b) SEM image of a WT gemma, c) SEM image of a 10-day-old WT thallus. d-i) Mp*dcl1a* mutant gemmae exhibiting the formation of supernumerary papillae and ectopic gemmae from the dorsal and ventral epidermis of the gemma. d-e) Mp*dcl1a-1^ge^* gemmae. f) 10-day-old Mp*dcl1a-1^ge^* thallus, g-h) Mp*dcl1a-2^ge^* gemmae. i) 10-day-old Mp*dcl1a-2^ge^* thallus. j-k) Mp*dcl1a-3^ge^* gemmae. k) 10-day-old Mp*dcl1a-3^ge^* thallus. Ectopic gemmae and papillae are highlighted using green or blue false-coloring, respectively. Scale bars: a, d, g, j = 0.5 mm. b-c = 200 μm. h, l, k = 500 μm. i = 400 μm.

### Transcriptomic landscape of Mp*dcl1a-3^ge^* mutants

We performed RNA-seq analysis in wild-type and Mp*dcl1a-3^ge^* gemmae and identified differentially expressed genes (DEGs). From the 3,436 DEGs identified, 1,567 corresponded to genes with increased expression and 1,869 showed reduced expression in Mp*dcl1a-3^ge^* gemmae (Figure 4a and Tables S2-S3). Consistent with a major role for Mp*DCL1a* in the biogenesis of miRNAs, Mp*dcl1a-3^ge^*mutant gemmae exhibit upregulation of 62 miRNA-precursors including 3 precursors giving rise to 3 of the 7 miRNA families conserved across land plants (MpmiR160, MpmiR166, and MpmiR408) (Figure 4b) and upregulation of 61 miRNA targets (Figure S5) (Tables S10-S11). Stem-loop RT-PCR assays confirmed that the increased levels of pre-miRNAs observed in Mp*dcl1a-3^ge^* correlate with a reduced expression of mature miRNAs (Figure S6). Similar to other plant systems^22^, we observed downregulation of a subset of miRNA-precursors in Mp*dcl1a-3^ge^* (Figure 4b and Table S11). We identified 55 transcription factors (TFs) with increased expression and 15 with reduced expression in Mp*dcl1a-3^ge^* gemmae (Figure S5 and Table S12-S13). Seven of the TFs upregulated in Mp*dcl1a-3^ge^* are targets of miRNAs (Figure S5). Consistent with these observations, Mp*dcl1a-3^ge^* mutants exhibit phenotypes observed in miRNA loss-of-function mutants including disruption of the regulatory module MpFRH1-MpRSL1^23,24^ (Figure S7) and alterations in phase change and apical dominance reported for mutants of miR529c^25^ and miR11671^26^ (Figure S8).

**Figure 4.**
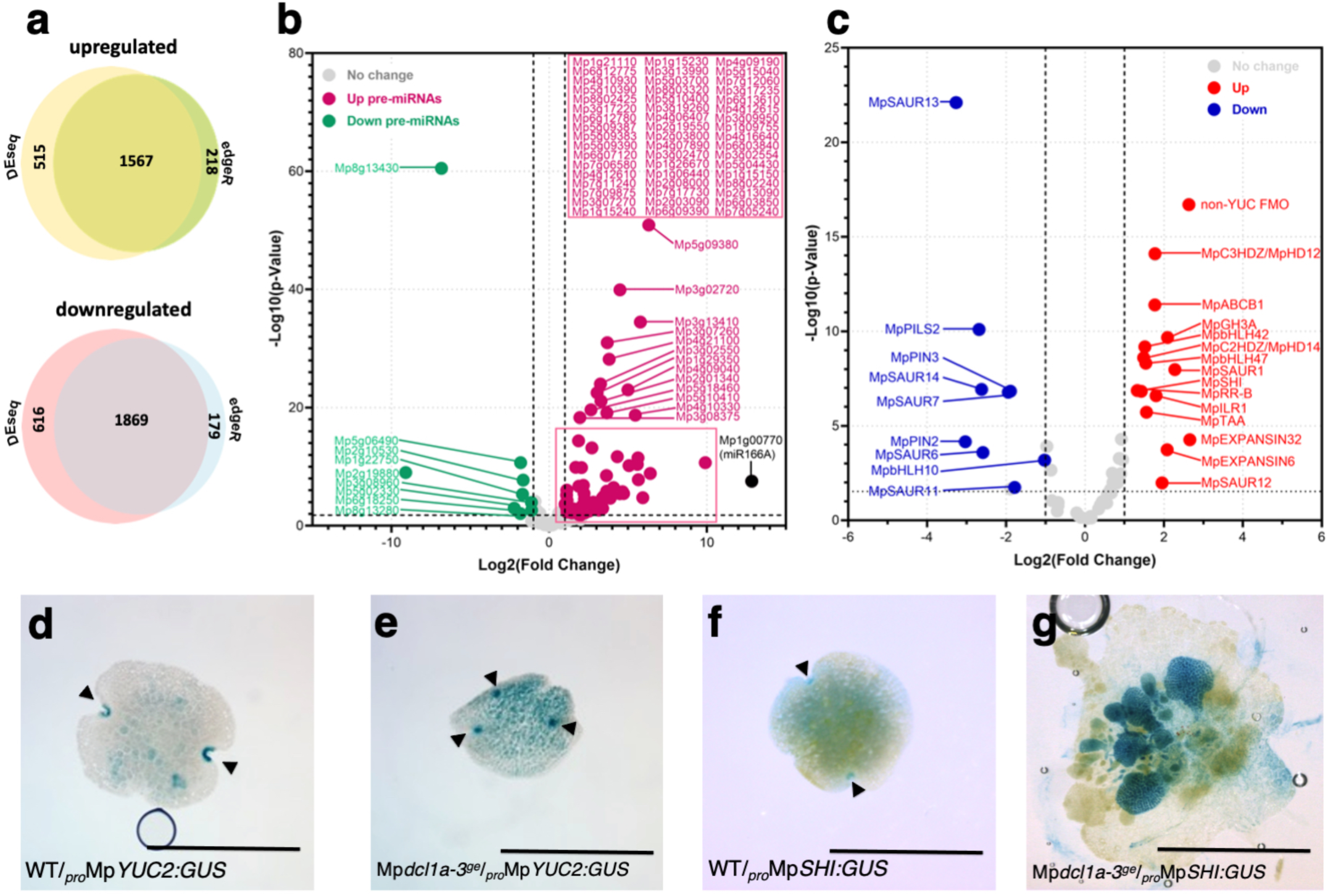
Transcriptomic analysis of Mp*dcl1a-3^ge^* gemmae. a) Identification of differentially expressed genes (DEGs) in Mp*dcl1a-3^ge^* gemmae shared between edgeR and DEseq2 analysis. b) Precursors of miRNAs (pre-miRNAs) differentially expressed in Mp*dcl1a-3^ge^* gemmae. The precursor of MIR166 (shown in black) is the third most highly expressed gene in Mp*dcl1a-3^ge^* gemmae. c) DEGs involved in auxin biosynthesis and signaling in Mp*dcl1a-3^ge^* gemmae. d) GUS staining observed in WT gemmae harboring the transcriptional fusion _pro_Mp*YUC2:GUS*. e) GUS staining observed in Mp*dcl1a-3^ge^* gemmae harboring the transcriptional fusion _pro_Mp*YUC2:GUS*. f) GUS staining observed in WT gemmae harboring the transcriptional fusion _pro_Mp*SHI:GUS*. g) GUS staining observed in Mp*dcl1a-3^ge^* gemmae harboring _pro_Mp*SHI:GUS*. Meristematic regions are indicated with black arrowheads. Scale bars = 0.5 mm.

We observed changes in basal levels of transcription of diverse genes involved in hormonal signaling pathways including auxin, ABA, cytokinin, ethylene, gibberellic acid and jasmonic acid (Tables S14-S19). Auxin signaling was particularly interesting since genes with increased expression included 4 of the 5 genes characterized so far that positively regulate tryptophan-dependent auxin biosynthesis including *TRYPTOPHAN AMINOTRANSFERASE OF ARABIDOPSIS 1* (Mp*TAA1*), Mp*SHI* (*in planta* confirmed in Figure 4g) and Mp*C3HDZ* ^21,27^ as well as a set of genes belonging to an auxin co-expression group including CLASS II *HOMEODOMAIN-LEUCINE ZIPPER* (Mp*C2HDZ*), *SMALL AUXIN-UP RNA1* (Mp*SAUR1*)*, BASIC HELIX-LOOP-HELIX42* (Mp*bHLH42*), *GRETCHEN HAGEN 3* (Mp*GH3*)^28^ and Mp*EXPANSIN6* ^29^ (Figure 4c and Table S14). We also identified 9 auxin-related genes with reduced expression in Mp*dcl1a-3^ge^* including Mp*SAUR6,* Mp*SAUR7,* Mp*SAUR11,* Mp*SAUR13*, Mp*SAUR14,* Np HLH10, *PIN-FORMED* (Mp*PIN*) and *PIN-LIKES* (Mp*PILS*) families (Figure 4c and Table S14). Collectively, these results support a role for Mp*DCL1a* as a major contributor to miRNA biogenesis and confirms its pleiotropic role in numerous hormone signaling pathways with a particular emphasis in the control of cell division and auxin signaling in gemmae.

### Formation of new auxin maxima territories in Mp*dcl1a-3^ge^*

Based on the expression pattern of Mp*YUC2* and Mp*SHI* ^18,21^, three major auxin maxima territories can be defined in wild-type gemmae, these are: apical notches, the cluster of rhizoid precursor cells and the stalk-anchoring cells^18^ (Figure 4d and 4f). Signal from both _pro_Mp*YUC2:GUS* and _pro_Mp*SHI*:*GUS* is enriched and expanded in the ectopic gemmae of Mp*dcl1a-3^ge^* relative to wild type (Figure 4e and 4g) which suggests the presence of novel sites of auxin biosynthesis and/or accumulation. Given that molecular tools to quantify auxin *in vivo* in *M. polymorpha* are yet to be developed, we established a protocol for immunolocalization of auxin using two commercially available antibodies (Materials and Methods). We observed clear and discrete auxin immunolocalization signals in the apical notches, rhizoid precursor cells and stalk-anchoring cells of wild-type gemmae (Figure 5a-d and S9). Treatment of wild-type gemmae with inhibitors of polar auxin transport (PAT): N-1-naphthylphthalamic acid (NPA) and 2,3,5-triiodobenzoic acid (TIBA) eliminated the auxin immunolocalization signal from rhizoid precursor cells (Figure S9b-c). A similar result was observed in wild-type gemmae treated with the inhibitors of auxin biosynthesis L-kynurenine and yucassin (Figure S9d-e). In Mp*dcl1a-3^ge^*, we observed increased auxin immunolocalization signal in the apical notches, the cluster of rhizoid precursor cells and ectopic gemmae (Figure 5e-h). Taken together, our results strongly suggest that Mp*dcl1a-3^ge^* exhibits altered auxin homeostasis that correlates with the formation of new auxin maxima territories in ectopic gemmae.

**Figure 5.**
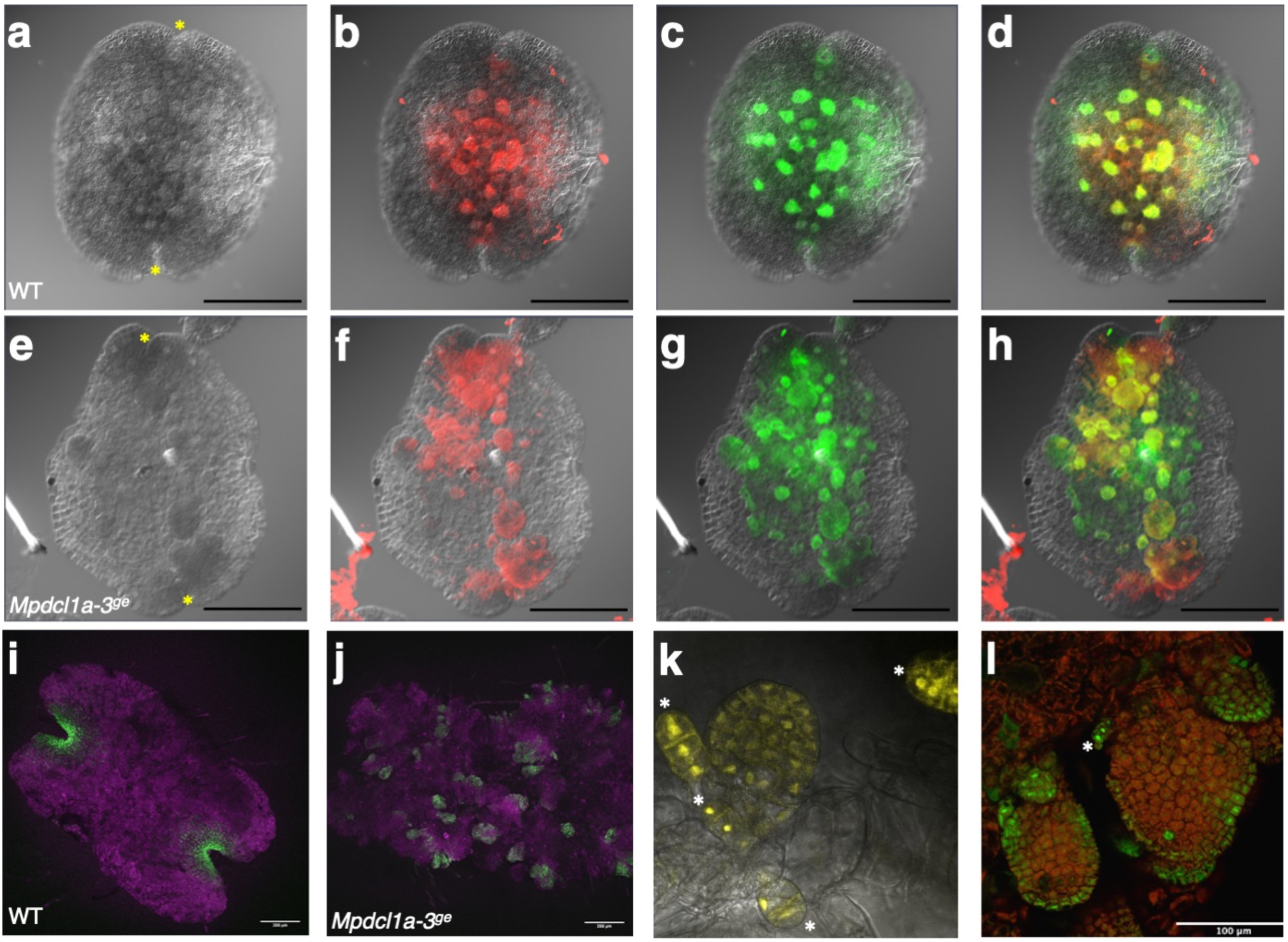
Formation of novel auxin maxima territories in Mp*dcl1a-3^ge^* gemmae. a) Differential Interference Contrast (DIC) microscopy image of a wild-type (WT) gemma, b-c) Double immunolocalization of auxin in WT gemmae using a monoclonal antibody that recognizes indoleacetic acid (IAA) (signal in red - panel b) and a polyclonal antibody that recognizes IAA (signal in green - panel c). d) Merged signals from panels b and c. e) DIC microscopy images of a Mp*dcl1a-3^ge^* gemma, f-g) Double Immunolocalization of auxin in Mp*dcl1a-3^ge^* gemmae h) Merged signals from panels f and g. Yellow asterisks indicate the location of apical meristems. i) EdU staining in WT gemmalings. j) EdU staining in a Mp*dcl1a-3^ge^* gemmalings. k) Expression pattern of the gemma-specific *gGROM:citrine* reporter in WT gemmae. l) Expression pattern of *gGROM:citrine* in Mp*dcl1a-3^ge^*. The presence of gemma initials is indicated with white asterisks. Scale bar = 1mm.

### Active DNA replication in Mp*dcl1a-3^ge^* ectopic gemmae

Given auxin stimulates the expression of genes involved in the regulation of cell cycle and DNA replication in a dose dependent manner^30,31^ and can promote cell division and organogenesis in the sites where it accumulates^32^, we hypothesized that the accumulation of auxin in ectopic gemmae promotes DNA synthesis and cell division. To test this, we performed staining with 5-ethynyl-20-deoxyuridine (EdU) to visualize DNA replication^33^. While in wild-type gemmalings the EdU signal was restricted to the two apical notches (Figure 5i), in Mp*dcl1a-3^ge^*gemmae the signal was highly enriched in ectopic gemmae (Figure 5j). These observations further support the role of Mp*DCL1a* as a regulator of patterning and cell division activity.

### The dorsal and ventral epidermis of Mp*dcl1a-3^ge^* acquires a gemminiferous cell fate

The developmental landscape of both the dorsal and ventral epidermis of Mp*dcl1a-3^ge^* gemmae resembles the floor epidermis of gemma cups that constitutes the gemminiferous region where gemmae are formed. To determine the identity of the ectopic tissues developing from the epidermis of Mp*dcl1a-3^ge^* gemmae, we employed a gemma marker named *gGROM:Citrine* (Ishizaki et al., unpublished) that is highly expressed in the epidermis of gemma initials and young gemmae, and is expressed at lower levels in mature gemmae (Figure 5k). In Mp*dcl1a-3^ge^*, the expression of *gGROM:Citrine* can be detected in the epidermis of gemmae, the ectopic elongated cells resembling gemma initials and also in young and mature ectopic gemmae (Figure 5l). We observed that the formation of ectopic gemmae in Mp*dcl1a-3^ge^* is 100% penetrant. (Figure S10a). Ectopic gemmae detached from the epidermis of Mp*dcl1a-3^ge^* plants develop into thalli that can form gemma cups and give rise to gemmae exhibiting the presence of ectopic gemmae (Figure S10b). Collectively, these observations indicate that the epidermis of Mp*dcl1a-3^ge^* gemmae recapitulates the morphology and the gemminiferous status of the single layer of the floor epidermis of gemma cups and demonstrate that ectopic gemmae that break the epidermis plane in Mp*dcl1a-3^ge^* acquire a gemma cell fate and this phenotype is 100% penetrant.

### Inactivation of auxin abolishes the formation of ectopic gemmae in Mp*dcl1a-3^ge^*

We employed a genetic approach to manipulate auxin levels in Mp*dcl1a-3^ge^* gemmae by expressing the bacterial gene *indoleacetic acid-lysine synthetase* (*iaaL*) capable of inactivating free auxin by conjugating free indole-3-acetic acid (IAA) with the amino acid lysine, under the constitutive promoter Mp*EF1* and using a β-estradiol inducible system (*pro*Mp*EF1:XVE)* as previously reported ^18^. The expression of *iaaL* results in decreased levels of free IAA in plant cells^34^ and the heterologous expression of *iaaL* in *M. polymorpha* leads to an auxin deficient phenotype characterized by the arrest of growth, the loss of all developmental patterns and the presence of undifferentiated cells^18^. We transformed spores obtained from the cross between Tak2 and Mp*dcl1a-3^ge^*, with the auxin conjugator *iaaL*, and randomly selected 31 plants for PCR genotyping. We screened for the presence of both the mutation in Mp*dcl1a-3^ge^* and the presence of the *iaaL* construct. 24 out of 31 plants that were genetically wild-type and positive for the presence of *iaaL* did not show the Mp*dcl1a-3^ge^* phenotype of ectopic gemmae. From the remaining 7 plants carrying the Mp*dcl1a-3^ge^* mutation and positive for the presence of *iaaL*, 4 displayed the phenotype of ectopic gemmae and 3 resembled the wild-type phenotype with flat gemmae (Figure S11). Given the mutant phenotype of ectopic gemmae in Mp*dcl1a-3^ge^* plants is 100% penetrant (Figure S10) and taking into account that in *M. polymorpha*, the β-estradiol inducible system can exhibit basal expression levels in the absence of the inducer ^35^, we wanted to test if the observed reversion of the mutant phenotype in the 3 lines correlated with the expression of *iaaL* (Figure S12). While all plants carrying the *iaaL* construct and grown for 7 days in media supplemented with 5 uM of β-estradiol under optimal conditions, exhibited characteristic auxin deficient phenotypes induced by *iaaL i.e.* arrest of growth, loss of all developmental patterns and the presence of undifferentiated cells ^18^ (Figure S13), revertant plants carrying the Mp*dcl1a-3^ge^* mutation but phenotypically wild-type (Figure S11) exhibited high levels of expression of *iaaL* in the absence of β-estradiol (Figure S12). Only one of the four genotypically and phenotypically Mp*dcl1a-3^ge^* mutant plants expressed *iaaL* but at lower levels than those observed in the revertant plants (Figure S12). Taken together, these results are consistent with an excess of auxin involved in the formation of ectopic gemmae.

### Disruption of the Mp*miR166a*-Mp*C3HDZ* module results in the formation of ectopic gemmae and alterations of auxin homeostasis

We hypothesized that the ectopic gemmae phenotype observed in Mp*dcl1a-3^ge^* mutants was due to the misregulation of either a single or multiple miRNA regulatory modules. Given that Mp*dcl1a* mutant gemmae showed defects in patterning and cell fate specification, they exhibited the formation of new auxin maxima territories and the precursor of miR166 is one of the top 10 genes with increased expression in Mp*dcl1a-3^ge^* (Figure 4b and Table S2), we focused on the MpmiR166a-Mp*C3HDZ* module involved in cell fate specification and control of auxin homeostasis in diverse land plants^27,36–39^. In angiosperms, miR166 acts as a mobile signal that promotes cell fate specification during embryonic and postembryonic development through post-transcriptional silencing of *C3HDZ* transcription factors, by restricting their spatial expression^36,40^. The genome of *M. polymorpha* encodes two copies of Mp*MIR166:* Mp*MIR166a* is expressed in gametophytic and sporophytic tissue while expression of Mp*MIR166b* is restricted to sporophytic tissue^10^. Mp*C3HDZ* is a confirmed target of miR166^5,10^. We generated loss-of-function alleles of Mp*MIR166A* by homologous recombination and gain-of-function alleles of Mp*C3HDZ* by expression of a Mp*miR166A*-resistant version of Mp*C3HDZ* driven by its own promoter (_pro_Mp*C3HDZ:*Mp*C3HDZ\**) or by the constitutive promoter EF1 (_pro_Mp*EF1:*Mp*C3HDZ\**). Gain-of-function alleles showed narrow thalli with reduced lateral expansion that produced gemma with ectopic gemmae (Figure 6b-c). Loss-of-function alleles of Mp*MIR166a* phenocopied the strong phenotype of Mp*C3HDZ* gain-of-function; mutant thalli were narrow without lateral expansion, branching was delayed and developed gemma with ectopic gemmae (Figure 6d). In Arabidopsis, *C3HDZ* transcription factors promote auxin biosynthesis by targeting the expression of *TAA* and *YUC* enzymes^27,39^. Collectively, these data suggest that transcriptional changes observed in Mp*dcl1a-3^ge^*mutants imply an excess of epidermal auxin synthesis and the formation of auxin maxima pools with morphogenetic effects that may be channeled through MpC3HDZ for the specification of gemma cell fate.

**Figure 6.**
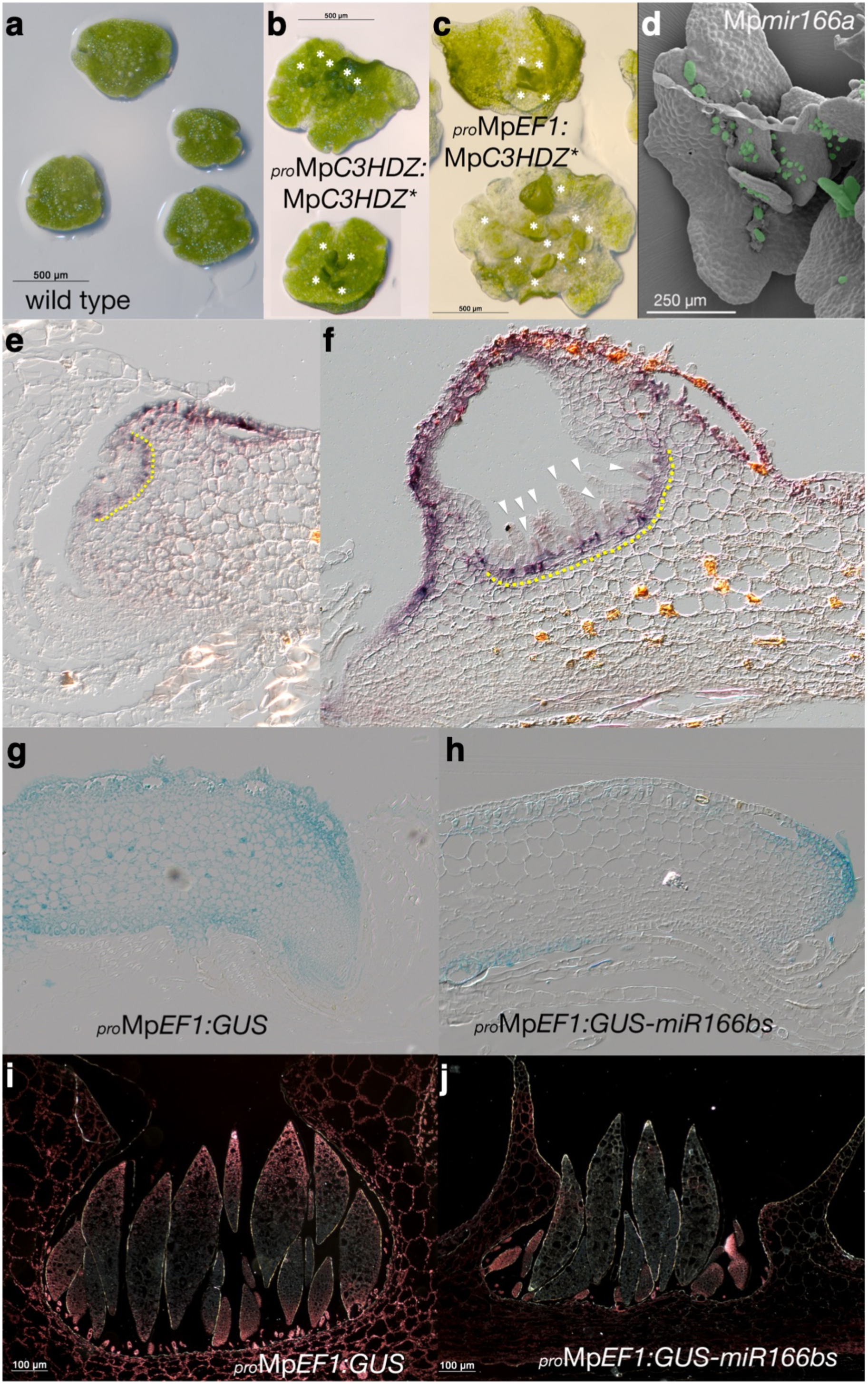
Disruption of the Mp*miR166a*-Mp*C3HDZ* regulatory module results in the formation of ectopic gemmae. a) WT gemmae. b) Gemmae from Mp*C3HDZ* gain-of-function plants expressing a miRNA-resistant version of Mp*C3HDZ*\* under its native promoter and exhibiting the presence of ectopic gemmae. c) Gemmae from Mp*C3HDZ* a gain-of-function plants expressing a miRNA-resistant version of Mp*C3HDZ*\* under the constitutive promoter Mp*EF1* exhibiting the presence of ectopic gemmae. White asterisks indicate the presence of ectopic gemmae. d) Gemmae from a *MIR166a* loss-of-function plant with ectopic gemmae. Ectopic gemmae are highlighted using green false-coloring. e) *In situ* hybridization of Mp*C3HDZ* showing a discrete signal in the dorsal epidermis of a young thallus and the base of a developing gemma cup. f) *In situ* hybridization of Mp*C3HDZ* showing a strong signal in the dorsal epidermis of a mature thallus and the base of the gemma cup and a weak signal in mature gemmae. The yellow dotted line indicates the floor epidermis of the gemma cup. White arrowheads indicate developing gemmae. g) Light field image shows broad and uniform GUS staining in the thallus of a line expressing GUS under the constitutive promoter Mp*EF1* (_pro_Mp*EF1:GUS*). h) Light field image of a thallus from plants expressing *GUS* harboring a *miR166* binding site (_pro_Mp*EF1:GUS-miR166bs*), staining is prominently reduced and just weak staining patches are observed in the apical lobe of thallus. i) Dark field image of gemmae from plants expressing _pro_Mp*EF1:GUS* showing GUS staining in gemmae at different developmental stages, the walls and floor of the gemma cup. j) Dark field image of gemmae from plants expressing _pro_Mp*EF1:GUS-miR166bs* showing GUS staining in young developing gemmae but staining is absent in mature gemmae and the walls and floor of the gemma cup.

### Mp*miR166*-mediated regulation of Mp*C3HDZ* is largely quantitative rather than qualitative

Mp*C3HDZ* is expressed in both the gametophyte and sporophyte generations based on RT-PCR analysis^38^. Mp*C3HDZ in situ* hybridization of sectioned mature gametophyte exhibited staining in the walls and basal cell layer of developing gemma cups but not in young developing gemmae (Figure 6e-f). To determine the expression territory of Mp*MIR166a* we compared the patterns reflected by *GUS* reporter genes with and without Mp*miR166a* binding sites driven by the Mp*EF1* promoter conferring constitutive expression (Figure 6g-h). A loss of reporter gene expression was evident in young differentiating ventral tissues as well as more differentiated dorsal tissues in lines harboring a Mp*miR166a* binding site (Figure 6h). In developing gemma cups, a reduction of *GUS* expression was observed in the walls and basal cell layer and developing gemmae (Figure 6i-j). The partial overlap in expression patterns of Mp*C3HDZ* and Mp*MIR166a* indicates that Mp*MIR166a* regulation of Mp*C3HDZ* in the *M. polymorpha* gametophyte is quantitative dorsally and qualitative ventrally.

## DISCUSSION

In this work we showed that the DCL family in *Marchantia polymorpha* is composed by five members: MpDCL1a, MpDCL1b, MpDCL3, MpDCL4 and MpmDCL, with MpDCL1b and MpmDCL encoding non-canonical DCLs. We found that Mp*DCL1a* is the major contributor for the biogenesis of miRNAs and its partial loss of function results not only in the accumulation of miRNA precursors and miRNA targets but in alterations in cell specification and patterning including the formation of ectopic gemmae from the epidermal cells of the gemma in a Class III HD-ZIP and miR166-dependent manner that correlates with the formation of additional auxin maxima territories. Collectively, these results demonstrate that DCL1, Class III HD-ZIP, and miR166 functioned in the development of the body of the last common ancestor of the vascular plants and bryophytes.

The partial loss-of-function of Mp*DCL1a* results in downregulation of a set of mature miRNAs, upregulation of both miRNA precursors and miRNA targets and the Mp*dcl1a-3^ge^* mutant also exhibit phenotypes of miRNAs loss-of-function mutants. This is consistent with that observed in *dcl1* mutants from other plant models positioned at different evolutionary nodes ^22,41–43^ and is in agreement with a major role for Mp*DCL1a* in the biogenesis of miRNAs in *M. polymorpha*. Transcriptomic analysis revealed the diversity of regulatory pathways and developmental programs influenced by miRNAs in *M. polymorpha* including diverse hormonal signaling pathways.

The phenotypical characterization of Mp*dcl1a* hypomorphic alleles revealed a potential homeotic change of rhizoid precursor cells into gemminiferous tissue. Despite the increased activity of rhizoid markers (_pro_Mp*FRH1:YFP*) in Mp*dcl1a* mutants, loss of MpDCL1a activity does not result in ectopic rhizoids such as those observed during pharmacological auxin treatment. This suggests that MpDCL1a and the genes targeted by MpDCL1a-dependent miRNAs can override the cell identity pathway established by the Mp*RSL1* gene. Ectopic gemmae have fully acquired gemma identity as shown by the gemma-specific marker *gGROM:Citrine* as well as their ability to develop mature thalli and complete sexual and asexual life cycles (in the case of Mp*dcl1a* weak alleles). Our transcriptomic and cytological data supports a role for Mp*DCL1a* as a repressor of auxin biosynthesis and cell-cycle. In Mp*dcl1a-3*^ge^ ectopic gemmae we observed the formation of new auxin maxima territories, active DNA replication and increased and expanded expression patterns of auxin biosynthesis related markers such as *pro*Mp*SHI*:*GUS* and *pro*Mp*YUCCA2:GUS*. This not only happens in the central rhizoid domain, but also in the apical notch itself, where increased auxin levels may be prematurely differentiating the Mp*dcl1a* thallus, causing delays in bifurcation events and a de-phasing of plastochron transitions. Furthermore, genetic evidence employing the auxin conjugator *iaaL* in the Mp*dcl1a-3*^ge^ background indicates that auxin plays a prominent role in the formation of ectopic gemmae. This result is consistent with the proposed role of auxin as a neutral organogenic currency, providing a momentum for differentiation without determining specific cell identities ^18,44^. Our study shows that Mp*DCL1a* and its dependent miRNAs control the availability of transcription factors (*e.g.* Mp*RSL* vs Mp*C3HDZ*) in specifying cell identity downstream of auxin in the rhizoid precursor cell domain during gemmae development.

Genetic and molecular evidence helped us uncover a main role for the Mp*MIR166a*-Mp*C3HDZ* regulatory module in the regulation of gemmae cell fate. In angiosperms, members of the *C3HDZ* family have multiple functions in pattering, polarity, vascular differentiation and shoot identity but it is unclear what is their shared role in bryophytes and tracheophytes^37,40,45–47^.

C3HDZ expression is posttranscriptional downregulated by miR165/6 across land plants while in angiosperms miR165/6 spatially restricts the expression of *C3HDZ* members in a qualitative fashion^37,45,48^. In Arabidopsis, *REVOLUTA* -a *C3HDZ* member-targets the promoters and activates the expression of *YUCCA5* and *TAA1*, and the gain-of-function of *REV* shows increased accumulation of auxin ^27,39^. Arabidopsis *dcl1-5* mutant embryos exhibit ectopic expression of *REV* and an increase in auxin signaling ^49^.

Our results show that *MIR166* loss-of-function mutants and Mp*C3HDZ* gain-of-function mutants exhibit the formation of ectopic gemmae suggesting a causal explanation for such phenotype observed in Mp*dcl1a* mutants. The partial loss-of-function of Mp*DCL1a* results in a reduction in the basal levels of Mp*MIR166a* with a concomitant increase in the transcript levels of its target Mp*C3HDZ* and possibly other targets (e.g. Mp*YUC2* itself is a target of a Marchantia-specific miRNA). Although speculative at present, we propose that increased levels of Mp*C3HDZ* results in: i) the increased expression of Mp*YUC2,* Mp*SHI*, Mp*TAA* and changes in gene expression of a battery of hormone and auxin signaling genes that lead to increased biosynthesis of auxin and formation of new auxin maxima territories -influenced by the downregulation of Mp*PIN* and Mp*PILS*- and ii) the morphogenetic effects of auxin maxima are channeled through Mp*C3HDZ* to promote the specification of the gemmae cell fate in rhizoid precursor cells despite the presence of Mp*RSL* transcription factors, resulting in one of the first ever reported homeotic phenotypes in liverworts to date. Interestingly, the regulators of gemma development described to date Mp*KAR*^15^, Mp*GCAM1*^16^ and Mp*KAI2*^50^ did not exhibit differential expression in Mp*dcl1a-_3ge_*.

Collectively, our results show that in wild-type gemmae, repression of Mp*C3HDZ* by Mp*MIR166a* suppresses gemmae cell fate and reinforces the observation that the gemmae developmental program shares a degree of deep homology with the axillary meristem developmental program in angiosperms ^17^, both of which are dependent on C3HDZ activity. Our data indicates that DCL1, Class III HD-ZIP, and miR166 functioned in the development of the body of the last common ancestor of the bryophytes and vascular plants and provides a novel working framework to interrogate the basic principles of cell specification and patterning in plants.

## MATERIALS AND METHODS

### Plant material and growth conditions

*M. polymorpha* Takaragaike-1 (Tak-1) (male) and Tak-2 (female) plants, were maintained asexually by *in vitro* culture as described in Kubota et al., 2013. Gemmae were plated on half strength Gamborg’s B5 medium (PhytotechLab) containing 1% agar and 1% sucrose. Plants were grown under continuous white fluorescent light (50–60 μmol photons m-2 s-1) at 22°C (Kato *et al*., 2015). After 4 weeks of *in vitro* culture, thalli were transferred to pots with rock wool and placed under white fluorescent continuous light supplemented with far-red light to induce the development of gametophores as previously reported (Chiyoda *et al*., 2008). To perform crosses, a drop of sterile water was placed on the antheridiophore until sperm was released, then the drop was collected and placed on top of archegoniophores (https://www.youtube.com/watch?v=YFjfgr-wsy0). For genetic transformation, we followed the protocol described in Sauret-Güeto et al., 2020 and employed spores obtained by crossing either Tak2 X Tak1 or Tak2 X Mp*dcl1a*, when necessary.

### Identification of DICER family members and phylogenetic analysis

For the identification of DICER proteins in *M. polymorpha* we obtained seed alignments from Pfam for all the characteristic domains and employed HMMER (ver 3.2.1) to interrogate the translated reference transcriptome (obtained from marchantia.info). We also identified DICER sequences by reciprocal best hit BLAST (E-value threshold = -10) using *A. thaliana* DICER proteins as a reference. For phylogenomic analysis, *DICER-LIKE* protein sequences were retrieved from published literature (Mukherjee *et al*., 2013; Coruh *et al*., 2015; Huang *et al*., 2015; You *et al*., 2017). Additional sequences were identified from Phytozome (ver 12) with BLASTN and TBLASTN using Arabidopsis’s *DICER-LIKE* sequences and an E-value threshold of -10. In cases of incomplete gene model predictions, FGENESH+ (Softberry Inc. New York, NY, USA) was used to predict coding sequences using Arabidopsis DICER-LIKE sequences as homology template. Supplemental Table 1 lists the species included in this study. Domains were annotated using information from the above publications and confirmed using Pfam Scan (https://pfam.xfam.org/search#tabview=tab0). Full-length coding sequences were aligned by translation with MUSCLE in Geneious version 6.1.8 (Kearse *et al*., 2012). Manual curation of the alignment was performed where necessary, including trimming edges of shorter transcripts to avoid spurious alignment. Codons containing >75% gaps in the final alignment were stripped before tree-building. Maximum likelihood phylogenetic trees were inferred by RAxML version 7.2.8 (Stamatakis *et al*., 2014) using a General Time Reversible (GTR) model with gamma distribution rate of heterogeneity. Support values are based on 100 bootstrap replicates. Branches with less than 95% support, were collapsed.

### Generation of Mp*dcl1a* mutants and description of plasmids

The names of primers and sequences are listed in Table S20. To generate the transcriptional fusion *_pro_*Mp*DCL1a*:*GUS*, a 5 kb region upstream of the start codon including the 5’UTR was amplified from genomic DNA using primers proMpD1a221F and proMpD1a221R and cloned into pDONR221 using BP Clonase II (Invitrogen). The resultant fragment was subcloned into pMpGWB304 (Addgene ID: 68632) (Ishizaki et al, 2015) with LR Clonase II (Invitrogen) to transcriptionally fuse it with *GUS* reporter gene. For genome editing of Mp*DCL1a* (Mp7g12090.1), guide RNAs (gRNAs) were designed to target the first exon of Mp*DCL1a*, we use the CasFinder tool (https://marchantia.info/tools/casfinder/) with the genome of *M. polymorpha* (version 3) to screen the first exon of *MpDCL1a* for candidates gRNAs, the search criteria was: length of 20 nt, a seed region of 8 nt and NGG as PAM sequence. We selected two gRNAs targeting the first exon. For the construction of the double stranded guide RNA, two complementary oligonucleotides were hybridized (Table S20) and inserted by InFusion cloning, into a pENTR vector harboring the promoter of the small RNA nuclear U6 of *M. polymorpha* (*pro*Mp*U6*) that drives the expression of each gRNA and the gRNA scaffold. The gRNAs were first cloned into a pENTR vector (pMpGE_En01) (Addgene ID:71534) and then subcloned by Gateway cloning into the binary vector (pMpGE010) (Addgene ID:71536) containing the *Cas9* gene driven by the promoter of the *M. polymorpha ELONGATION FACTOR1 alpha* (*pro*Mp*EF1*).

### miRNA-resistant Mp*C3HDZ* vector construction

Primers were designed to replace the wild-type sequence encoding the miR166 binding site in an MpC3HDZ cDNA. Three silent changes were introduced towards the 3’ end of the miRNA binding site as changes in this region have been shown to disrupt miRNA binding (Mallory et al., 2004).

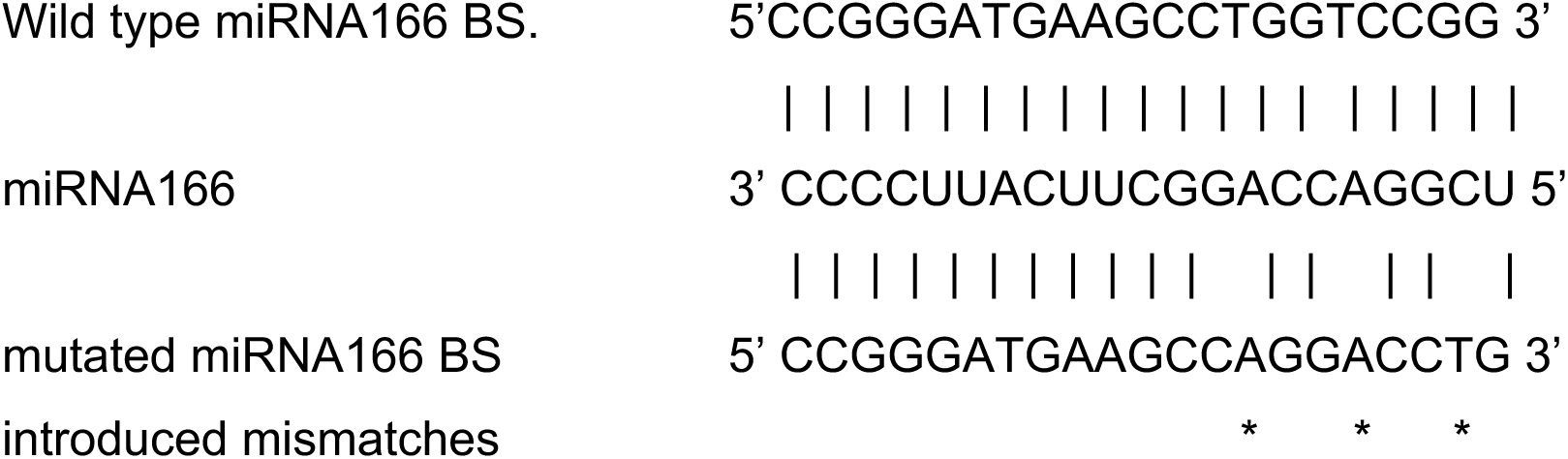

### Gene targeting vector construction

The method for homologous recombination-mediated gene targeting described by Ishizaki et al. (Ishizaki et al., 2013) was used to target the Mp*MIR166A* gene. We modified the recombination vector by using In-Fusion (Clontech) to insert the Gateway L1-L2 site into pJHY-TMp1 linearized by digestion with PacI. 5’ and 3’ homologous arms were amplified by PCR from genomic DNA. The 5’ arm was amplified to TOPO clone it directly into the pENTRD-TOPO vector (Invitrogen), then recombined into the modified pJHY-TMp1 using Gateway LR Clonase II (Invitrogen). The 5’ arm was a 1.5 kb fragment and the 3’ homologous arm a 2.5kb fragment. The two fragments were 240 nt apart so that the recombination eliminates the Mp*MIR166A* stem-loop region.

### Plant transformation

Transformation of *M. polymorpha* with *Agrobacterium tumefaciens* was performed using 7-day-old sporelings (from a cross between Tak-2 X Tak-1 or Tak2 X Mp*dcl1a-3^ge^*) following the protocols described previously (Ishizaki et al., 2008; Ishizaki et al., 2013; Sauret-Güeto *et al*., 2020). The constructs for genome editing and transcriptional fusions were introduced into the *A. tumefaciens* strain GV2260. After transformation, transgenic plants were selected in half strength B5 Gamborg media supplemented with either hygromycin (10 μg/ul) or chlorsulfuron (0.5 mM).

### Screening and genotyping of mutant plants

Plants growing on selection media were screened by PCR genotyping, gRNA-targeted regions were amplified from genomic DNA isolated from the T1 plants with GoTaq Green Master Mix (Promega) using the primers g1-seqF/g1-seqR and g2-seqF/g2-seqR for gRNA1 and gRNA2, respectively. PCR products were purified and sequenced by Sanger sequencing with g1-seqF and g2-seqF, respectively. Homologous recombination transformants were also screened by PCR to verify absence of the wild-type gene. Primers flanking the region targeted for replacement were designed and used for Terra Direct genotyping PCR (Clontech) to identify transformants lacking the wild-type fragment. Because the insertion fragment length exceeds the maximum size for the Terra direct polymerase, the PCR was repeated for plants for which the wild-type fragment failed to amplify in the Terra Direct PCR. After allowing individual plants to grow for three more weeks, DNA was extracted from potential homologous recombination transformants using the DNeasy kit (Qiagen) and standard PCR was performed using ExTaq HS kit (TaKaRa). Control reactions were run using wild-type DNA as a template.

### Segregation of CRISPR transgene

The mutant alleles Mp*dcl1a-2^ge^* and Mp*dcl1a-3^ge^*were crossed with wild-type plants to segregate the CRISPR-Cas9 machinery and generate male and female individuals for each line. PCR amplification products obtained from Mp*dcl1a-2^ge^* and Mp*dcl1a-3^ge^* mutant alleles are reduced in length relative to non-edited wild-type plants. Plants with edition at the target site and without Cas9 were selected by PCR for further analysis. For the inducible *iaaL* x Mp*dcl1a-3^ge^* experiment we transformed spores obtained from the cross between Tak2 and Mp*dcl1a-3^ge^*, with the auxin conjugator *iaaL*, and randomly selected 31 plants for PCR genotyping. We screened for the presence of both the mutation in Mp*dcl1a-3^ge^* and the presence of the *iaaL* construct by PCR. Primer sequences are listed in Table S20.

### RNA isolation

Total RNA was extracted from wild-type, Mp*dcl1a-1^ge^*and Mp*dcl1a-3^ge^* gemmae using Trizol Reagent (Thermo Fisher Scientific) following the manufacturer’s protocol. Three biological replicates were extracted from each genotype. Each replicate consisted of RNA from around 60 gemmae freshly collected from gemma cups. The quality and quantity of the total RNA were evaluated using a NanoDrop 2000 spectrophotometer (Thermo Fisher Scientific) and a Bioanalyzer 2100 (Agilent).

### GUS histochemical assay

To determine the activity of the enzyme beta-glucuronidase (GUS) in transgenic plants, a substrate buffer was used for the enzyme: 10 mM EDTA, 0.1% Triton X-100, 0.5 mM potassium ferricyanide, 0.5 mM ferrocyanide of potassium, 100 μg / mL of chloramphenicol, 1 mg / mL of 5-bromo-4-chloro-3-indolyl-BD-glucuronic acid (X-gluc) in 50 mM of phosphate buffer. Plants were incubated in the GUS staining buffer at 37°C overnight and then were cleared with ethanol.

### Auxin immunolocalization

Gemmae were fixed with 2.5% PFM in 1X PBS plus 2% Tween 20 at 4°C overnight then washed three times with 1X PBS plus 0.2% Tween 20 on ice. For disruption of cell walls, samples were placed in an enzymatic digestion solution: 1% driselase (D8037), 0.5% cellulase (C1794), 1% pectolyase (P5936), 1% BSA (all reagents from Sigma) at 37°C for 1 hour and washed with 1X PBS plus 0.2% Tween 20 for 15 minutes. For permeabilization, samples were incubated with 1X PBS plus 2% Tween 20 for 2 hours, on ice. For labeling, samples were incubated with both the primary polyclonal anti-IAA antibody (Agrisera, AS09 421) and the primary anti-auxin monoclonal antibody (SIGMA A0855) at a dilution of 1:500 overnight in an horizontal shaker at 4°C, washed and labeled with either the secondary anti-rabbit antibody conjugated to red fluorescent dye CF^TM^555 at dilution of 1:300 or the secondary anti-mouse antibody conjugated to green fluorescent dye CF^TM^488A at dilution of 1:200, overnight in an horizontal shaker at 4°C. Samples were mounted in a coverslip with Prolong-Gold antifade mountant reagent.

### Pharmacological assays

Assays were designed as follows: two replicates with 30 gemmae each. Gemmae were incubated in liquid Gamborg media supplemented with each inhibitor and placed on a shaker for 24 h. The pharmacological treatments were NPA 50uM, TIBA 100uM, L-kynurenine 100uM and yucassin 100uM. Gemmae were fixed to perform auxin immunolocalizations.

### EdU uptake assays

1-day-old gemmalings were incubated in liquid ½ Gamborg supplemented with 10 μM EdU (Click-iT™ EdU Cell Proliferation Kit for Imaging, Alexa Fluor™ 555 dye) overnight at 22°C under continuous white light in a shaker. After EdU uptake, gemmalings were fixed with 3.7% paraformaldehyde in 1X PBS for 30 min under vacuum and then washed two times with 3% BSA in 1X PBS. For tissue permeabilization, samples were placed in a solution of 0.5% TritonX-100 in 1X PBS for 30 min and washed twice with 3% BSA in 1X PBS. Coupling of EdU to the Alexa Fluor substrate was performed according to the manufacturer’s instructions. Samples were mounted in a slide with Prolong-Gold (P36930 Thermofisher) antifade mountant reagent.

### *In situ* hybridization

*In situ* hybridizations were performed as previously described (Floyd and Bowman, 2006).

### Microscopy analysis

All the observations in bright field for phenotypic analysis were performed with a stereomicroscope (LEICA S8AP0). For confocal laser scanning microscopy (LEICA SP8), samples stained with Alexa 555–labeled EdU and red fluorescent dye CF^TM^555 (for auxin immunolocalization) were excited at 555 nm. For scanning electron microscopic analysis, gemmae were taken directly from the gemma cup and fixed with glutaraldehyde and osmium tetroxide (Tsuzuki et al., 2019), then the samples were dehydrated in ethanol and dried. The samples were coated with gold. Images were captured with a scanning electron microscope (FEI Quanta 250 FEG).

### Growth rate experiments assays

Wild-type and Mp*dcl1a-3^ge^* gemmae were grown on solid B5 medium. We recorded the growth of thirty-eight plants from gemmae to mature thallus during fourteen days and we took pictures every two days with a stereomicroscope (LEICA S8AP0). Growth was measured using color threshold settings in the Fiji software as described in Schindelin et al., 2012.

### RNA sequencing and analysis

RNA-seq experiments were performed from three biological replicates of wild-type (Tak-1) and Mp*dcl1a-3^ge^* gemmae. Total RNA was extracted with TRIzol (ThermoFisher Scientific, 15596026) according to the standard protocol and library preparation with TruSeq kit. Sequencing was done with Illumina NextSeq 500 1×75 single end. Raw sequence reads were trimmed with trimmomatic v.0.36 (Bolger et al., 2014) and quality control analysis with FastQC v.0.11.9 (Wingett and Andrews, 2018) and MultiQC v.1.11 (Ewels et al., 2016). We used *M. polymorpha* v.6.1 reference genome (marchantia.info/download/MpTak_v6.1/) for alignment with Bowtie2 v.2.4.5 (Langmead & Salzberg, 2012) using --sensitive option and reads counts were obtained with Salmon v.1.8.0 (Patro et al., 2017). DEGs were identified using edgeR v.3.38.2 (Robinson et al, 2009) with a threshold log2FoldChange +/- 1 and FDR <0.05, and DESeq2 v.1.36.0 (Love et al., 2014) with a log2FoldChange +/- 1 and an adjusted pvalue <0.05. The selected DEGs are the ones shared in both analyses. GO-term enrichment analysis was performed using PlantRegMap (http://plantregmap.gao-lab.org/go.php), g:profiler (https://biit.cs.ut.ee/gprofiler/) and REVIGO (http://revigo.irb.hr/). Plots were made with Graph Pad Prism v.9.3.1.

## ACKNOWLEDGEMENTS

We gratefully acknowledge the following institutions and agencies for their support: M.A.A-V [National Council of Science and Technology (CONACYT) Ciencia Básica grants CB-158550 and A1-S-38383]; A.E.D.A. (CONACYT Ciencias de Frontera grant 23134); M.A.A-V and A.E.D.A. (UCMEXUS-CONACYT collaborative grants 19941-44-OAC7, CN-20-166 and CN-1798); M.A.A-V and J.H. (The Royal Society - Newton Advanced Fellowship grant NA15018); K.I. (MEXT KAKENHI grant 17H06472); R.A.M. (National Science Foundation grants IOS-1546825 and IOS-2247914; J.T.T. (National Institutes of Health grant T32-GM008659). L.D. (Austrian Academy of Sciences); D.G. and M.A.A-V. (Agropolis Fondation grant 1502-307) J.H. (BBSRC/EPSRC OpenPlant Synthetic Biology Research Centre grant BB/L014130/1); E.F.S and J.L.B (Australian Research Council grant DP200100225 and Australian Research Council Centre of Excellence for Plant Success in Nature and Agriculture grant CE200100015). We would like to thank Greta Hanako Rosas Saito for her assistance with the generation of SEM images. A.A.C. was the recipient of a scholarship (629316) from CONACYT.

## SUPPLEMENTARY INFORMATION

**Supplementary Figure 1.**
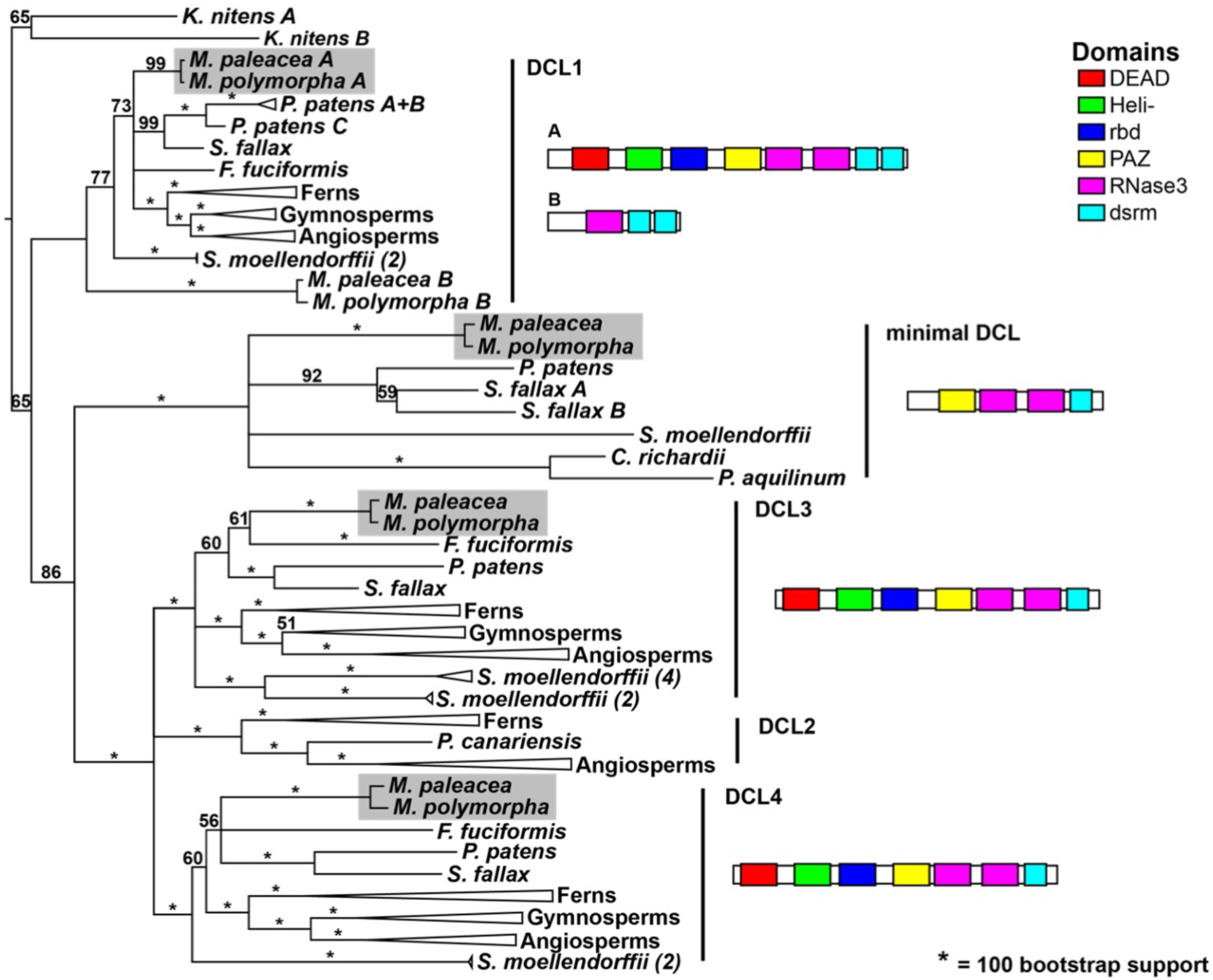
Phylogenetic reconstruction of the DICER-LIKE family in *M. polymorpha*. We identified five *DICER-LIKE* (*DCL*) genes in *M. polymorpha*. The evolutionary relationship of this family of genes was inferred through maximum-likelihood based phylogenetic analysis. Marchantia species possess a distinct DCL1 paralog and a minimal DCL in addition to canonical DCL3 and DCL4. Bootstrap support values are shown. Domain structure of Marchantia DCL paralogs is shown on the right. A Maximum Likelihood tree obtained with RAxML, based on an alignment of full-length coding sequences with codons containing >75% gaps stripped is shown. Numbers in parentheses at the end of each species corresponds to the number of orthologs in collapsed branches. Species in each group for other collapsed branches are listed in Table S1.

**Supplementary Figure 2.**
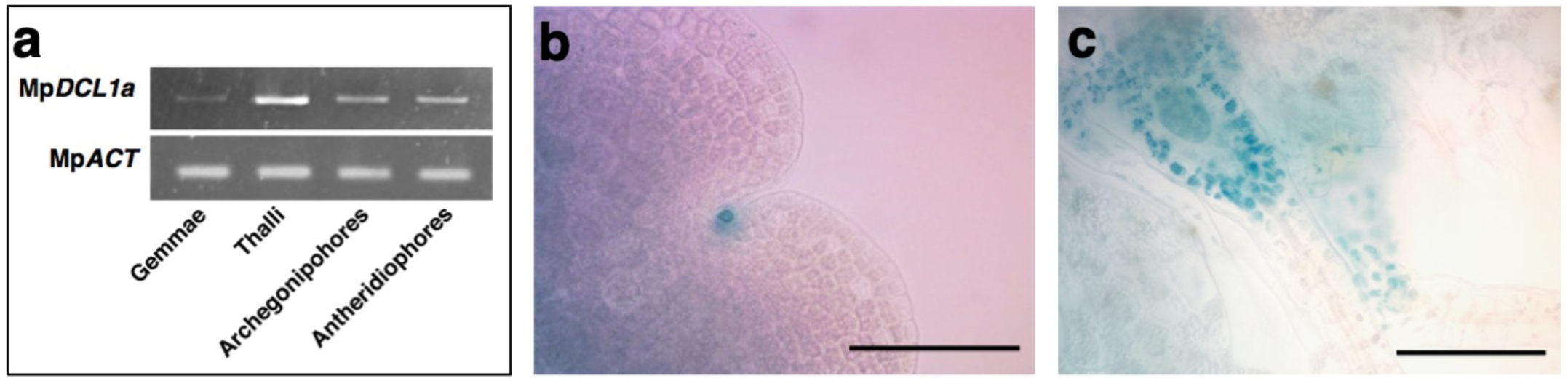
Expression profile of Mp*DCL1a*. a) Semi-quantitative RT-PCR analysis of Mp*DCL1a* in gametophytic and reproductive tissues, we used Mp*ACTIN7* as a loading control. b-c) Complementary expression pattern of Mp*DCL1a* obtained using the transcript ional fusion *pro*Mp*DCL1a*:*GUS* in a) papillae, b) archegonia. Scale bars = 0.1 mm.

**Supplementary Figure 3.**
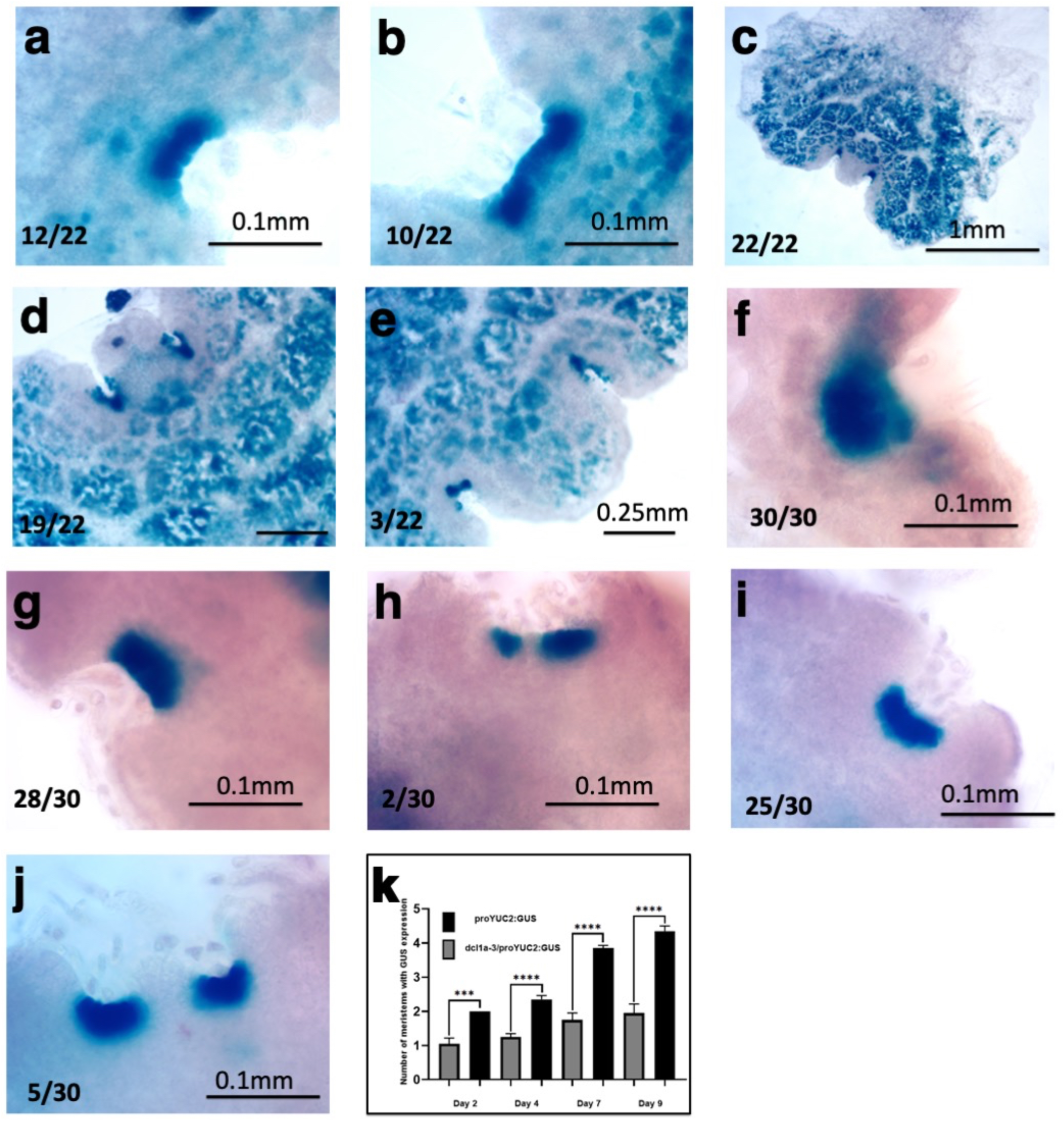
The timing of meristem division is affected in Mp*dcl1a-3^ge^* mutant plants. Developmental series of meristem divisions in wild-type and Mp*dcl1a-3^ge^* plants expressing the apical marker *pro*Mp*YUCCA2:GUS*. Wild-type plants show a regular division pattern, but division of the meristem is delayed in Mp*dcl1a-3^ge^*plants. Quantification of phenotypes are shown at the left bottom corner on each panel. a) 12 out of 22 4-days-old wild type plants show expansion of the meristematic zone. b) 10 out of 22 4-days-old wild type plants begins the first division of the meristem. c) After 7 days of development all wild-type plants analyzed show a fully divided meristem and complete differentiation of the central lobe. d) After 9 days of development the central lobe continues growing and the second division of the meristem begins in some plants (e) (3 out of 22). f) All M*pdcl1a* plants show expansion of the meristem after 4 days of development. g) The great majority of the Mp*dcl1a-3^ge^* plants (28 out of 30) exhibit an increase in the density of cells with meristem identity without division after 7 days of development. h) A scarce number of Mp*dcl1a-3^ge^* plants (2 out of 30) begin the first division of the meristem. i) The density of the meristematic zone continues increasing after 9 days without development in most of the Mp*dcl1a-3^ge^* plants (25 out of 30). j) The expansion of the central lobe begins in some Mp*dcl1a-3^ge^* plants after 9 days. k) Quantification of apical notches in wild-type and Mp*dcl1a-3^ge^* plants. Asterisks (****p < 0.0001, *** p< 0.005) indicate differences statistically significant in the number of meristems displaying GUS expression.

**Supplementary Figure 4.**
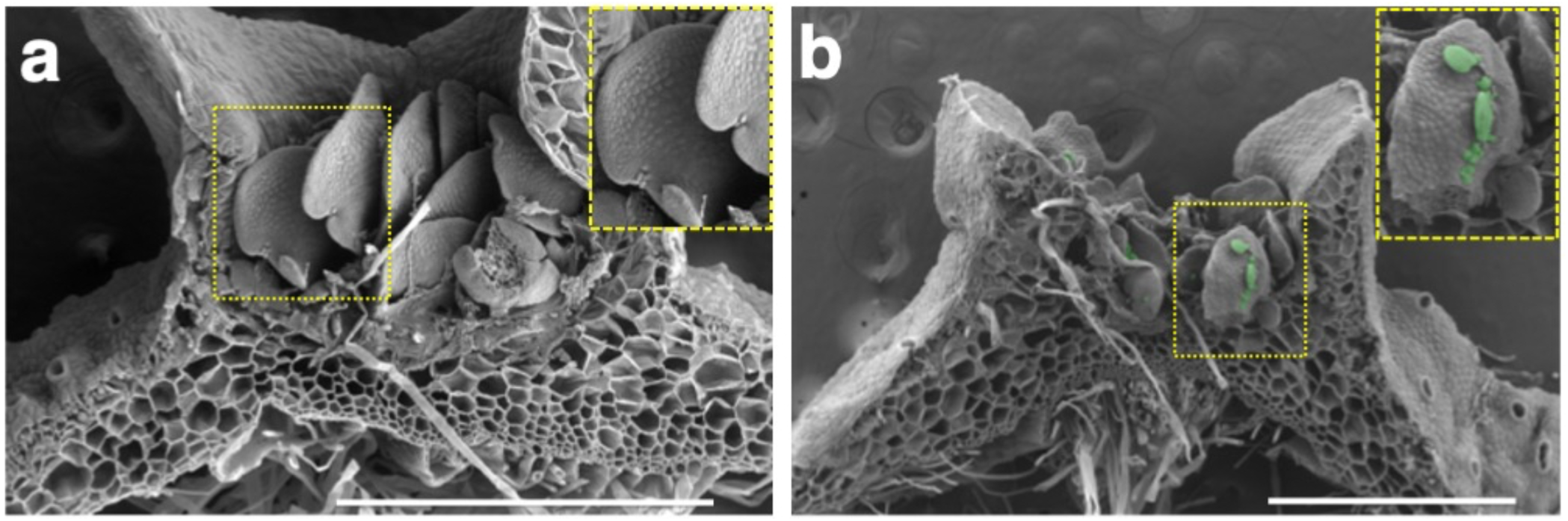
The epidermis of Mp*dcl1a* gemmae resembles the floor of gemma-cup. a) SEM image from a longitudinal section of a WT gemma cup. A magnification of the WT gemma is shown in the inset. b) SEM image from a longitudinal section of a Mp*dcl1a*-*3^ge^* gemma. A magnification of the mutant gemma exhibiting ectopic gemmae is shown in the inset. Ectopic gemmae are highlighted using green false coloring. Scale bar = 1mm.

**Supplementary Figure 5.**
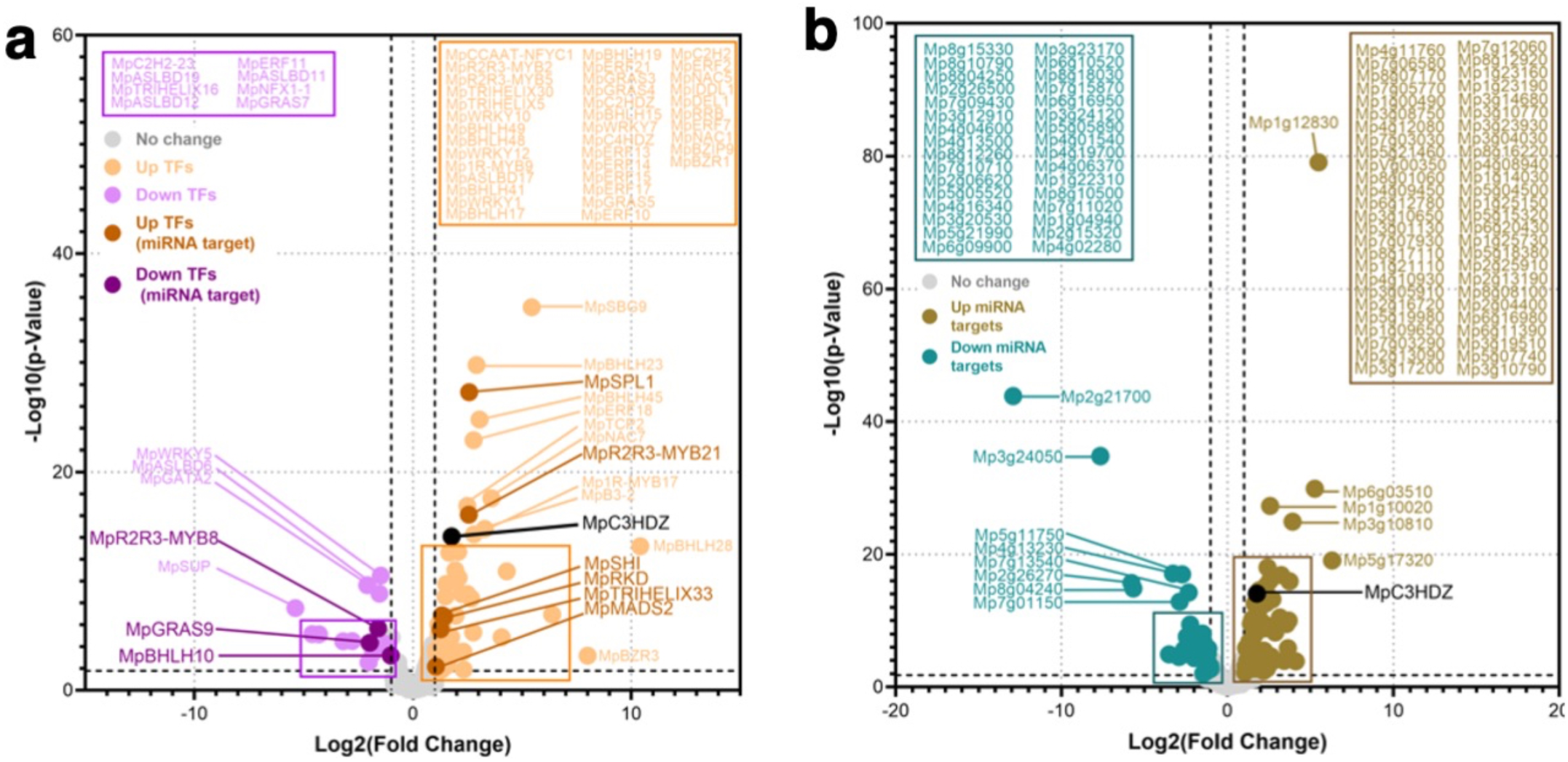
Transcription factors (TFs) and targets of miRNAs differentially expressed in Mp*dcl1a-3^ge^* gemmae. a) TFs differentially expressed in Mp*dcl1a*-*3^ge^* gemmae. The graph shows statistically significant (log2 fold change +/- 1 and FDR < 0.05) DEGs exhibiting increased (orange) or reduced (purple) levels in Mp*dcl1a*-*3^ge^*gemmae. Non-differentially expressed genes are depicted as gray dots. b) Targets of miRNAs differentially expressed in Mp*dcl1a*-*3^ge^* gemmae. The graph shows statistically significant (log2 fold change +/- 1 and FDR < 0.05) DEGs exhibiting increased (green) or reduced (blue) levels in Mp*dcl1a*-*3^ge^* gemmae. Non-differentially expressed genes are depicted as gray dots. TFs reported as targets of miRNAs exhibiting increased or reduced expression in Mp*dcl1a*-*3^ge^* gemmae are highlighted in dark orange and dark purple, respectively. Mp*C3HDZ* is targeted by miR166 and is shown in black bold text.

**Supplementary Figure 6.**
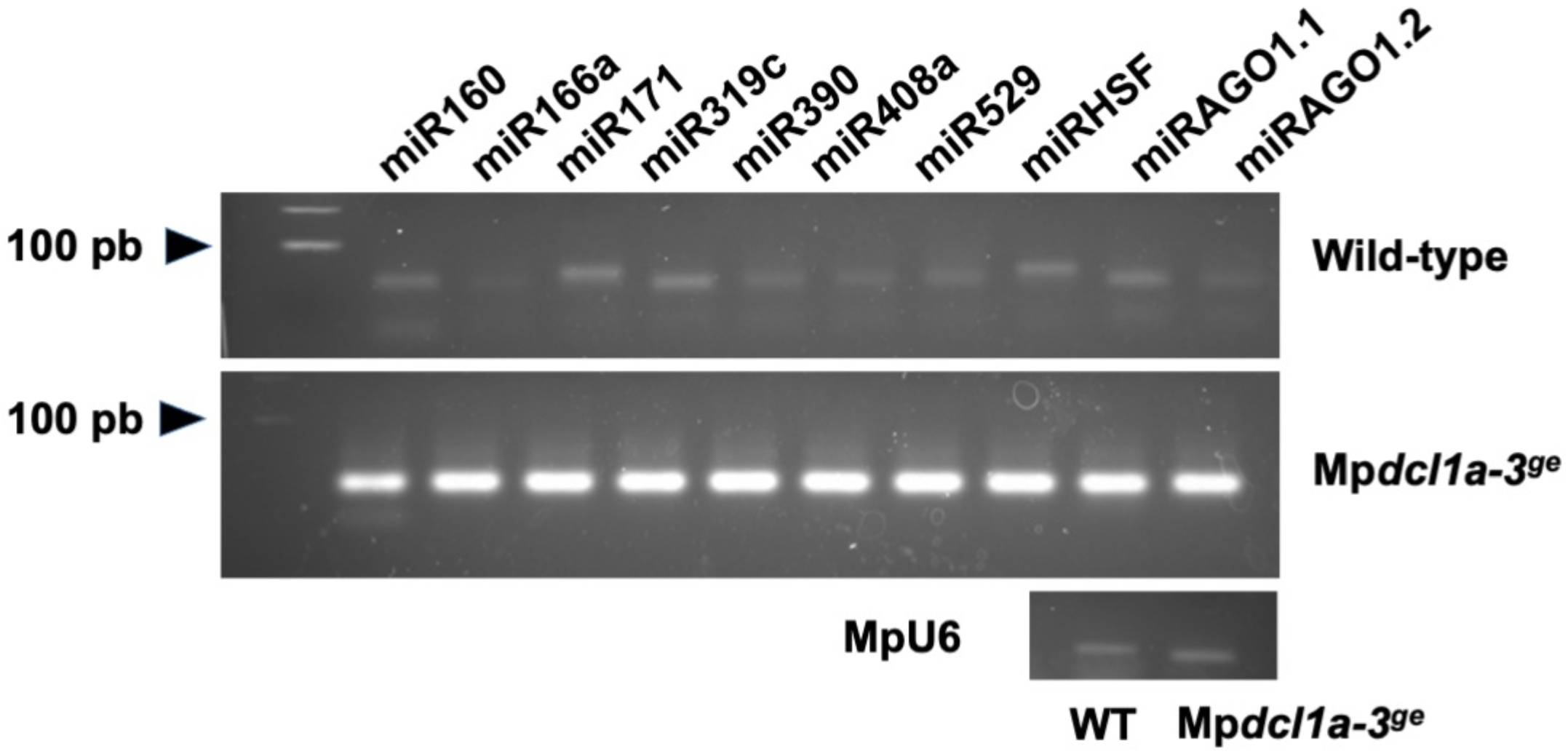
miRNA levels in Mp*dcl1a-3^ge^* are dramatically reduced. Stem-loop end-point RT-PCR assays from 14-days-old wild-type and Mp*dcl1a-3^ge^* plants. MpU6 was used as a loading control. In wild-type gemmae miRNA amplicons (60-70pb) are shown under 100 bp, in Mp*dcl1a-3^ge^* gemmae there are not miRNA amplicons, the bands shown are primer dimers.

**Supplementary Figure 7.**
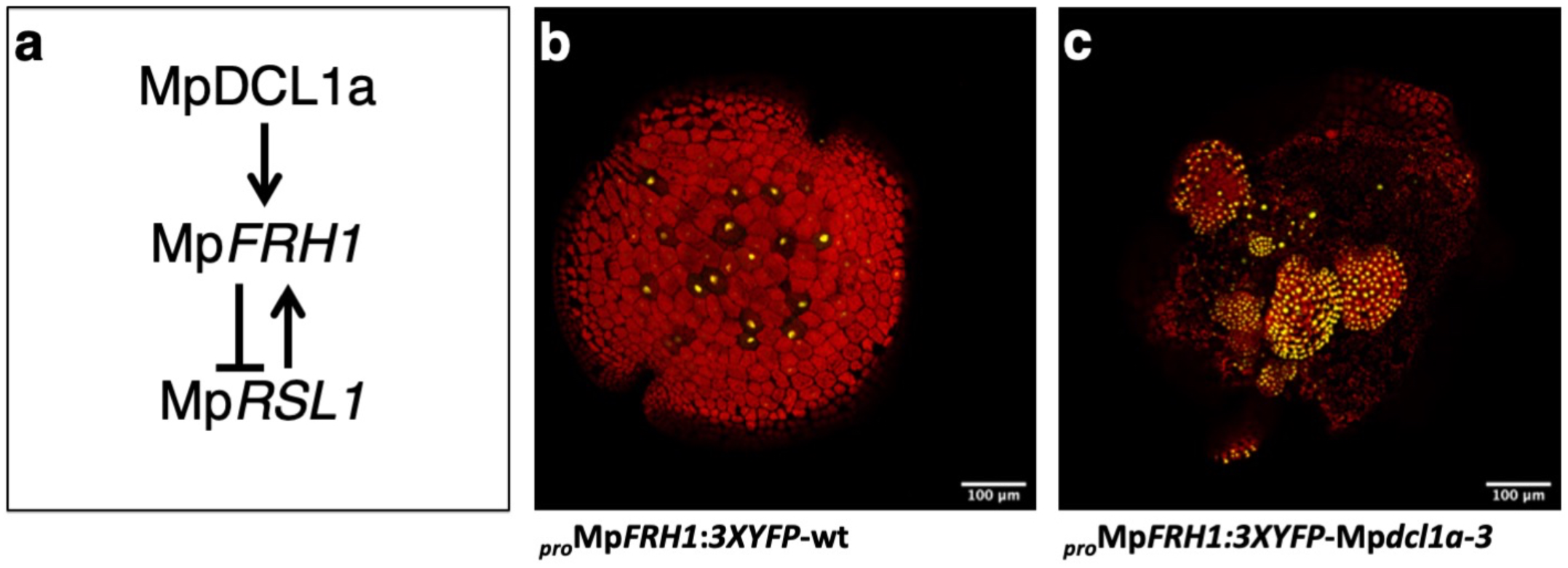
The negative regulation imposed by the miRNA Mp*FRH1* over Mp*RSL1* is released in Mp*dcl1a-3^ge^* mutants. a) MpRSL1 promotes the expression of Mp*FRH1* and in turn it is postranscriptionally regulated by Mp*FRH1*. b) Expression of the rhizoid marker *pro*Mp*FRH1:3XYFP* consisting of the promoter of the miRNA Mp*FRH1* driving the expresion of the yellow fluorescent protein (YFP) in a wild-type gemma (Honkanen, 2018). The expression is restricted to rhizoid precursor cells located in the central region of the gemma. b) Expression of *pro*Mp*FRH1:3XYFP* in Mp*dcl1a-3^ge^* gemmae bearing ectopic gemmae. The YFP signal can be observed in most of the epidermis of ectopic gemma at different developmental stages and it is now bearly detectable in the “bearing” gemma. In the Mp*dcl1a-3^ge^* mutant background MpRSL1 is released from the negative regulation of Mp*FRH1* and it results in the expansion of the expression territory of Mp*FRH1*.

**Supplementary Figure 8.**
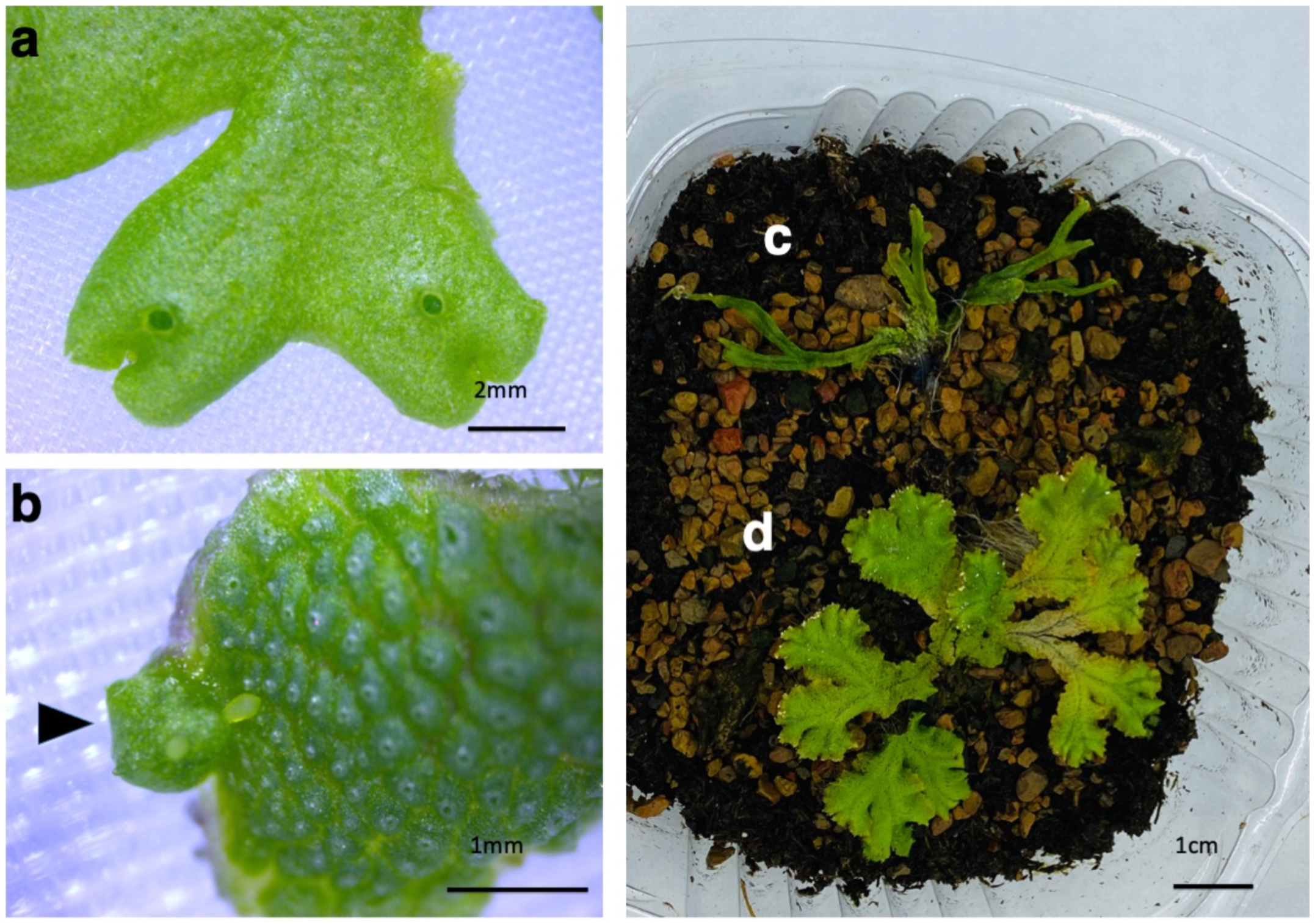
Mp*dcl1a-3^ge^ mutant* plants recapitulate the phase change and apical dominance phenotype observed in single miR529c and miR11671 mutants. a) 21-days-old wild-type thallus grown under continuous white light. b) A developing antheridiophore in a 21-days-old Mp*dcl1a-3^ge^* thallus grown under white continuous light is shown (black arrowhead). The phenotype observed in Mp*dcl1a-3^ge^* phenocopies the phenotype observed in mir529c loss-of-function mutants (Tsuzuki, 2019). c) 4-weeks-old Mp*dcl1a-3^ge^* plant growing in soil. d) 4-weeks-old wild-type plant growing in soil. The phenotype of apical dominance observed in Mp*dcl1a-3^ge^* phenocopies the phenotype observed in miR11671 loss-of-function mutants (Streubel et al., 2023).

**Supplementary Figure 9.**
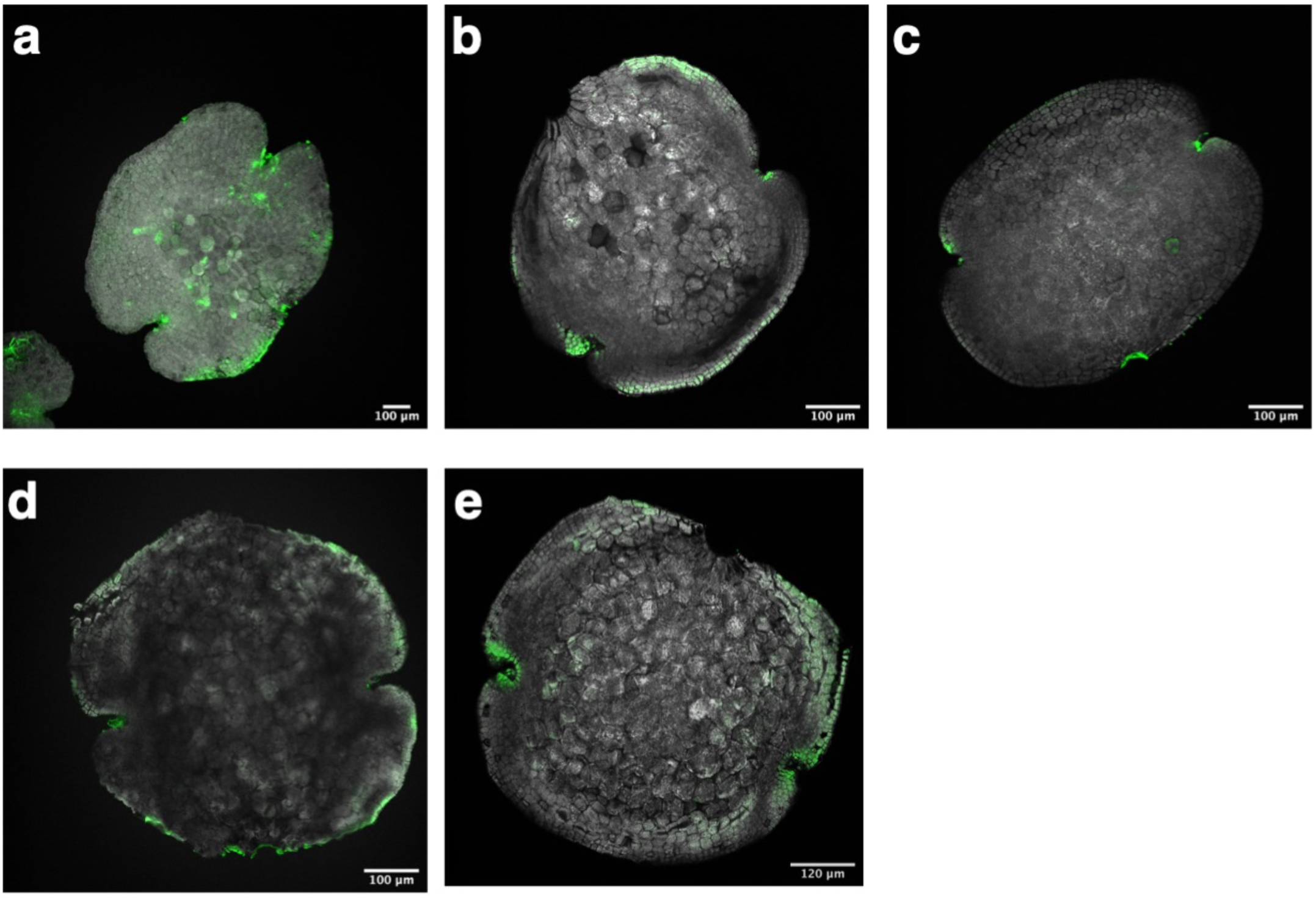
Immunolocalization of auxin in plants treated with inhibitors of either the biosynthesis or transport of auxin. a) Discrete auxin immunolocalization signals from mock treated wild-type gemmae can be observed in the apical notches, the precursors of rhizoids, rhizoids and the cells formerly attaching the gemma to the stalk. b) Auxin immunolocalization signals from gemmae treated with NPA (50 uM). c) Auxin immunolocalization signals from gemmae treated with TIBA (100 uM). d) Auxin immunolocalization signals from gemmae treated with L-kynurenin (100 uM). e) Auxin immunolocalization signals from gemmae treated with yucasin (100 uM). In wild-type gemma, auxin maxima territories can be observed in the apical notches, the precursors of rhizoids, rhizoids and the cells formerly attaching the gemma to the stalk (Figure 7a-d). The immunolocalization pattern of auxin maxima territories overlaps with the expression pattern of the auxin biosynthesis genes and components of the nuclear auxin signaling pathway (Eklund et al., 2015; Flores-Sandoval, 2015; Figure 6e and 6g). The immunolocalization pattern of auxin maxima territories is altered in wild-type gemmae treated with the inhibitors of Polar Auxin Transport (PAT): NPA and TIBA, auxin maxima is restricted to apical notches and the cells formerly attaching the gemma to the stalk and are no longer detectable in the rhizoids precursors cells. In gemmae treated with inhibitors of biosynthesis (L-kynurenin and yucassin), the auxin maxima territories are similar to those observed in gemmae treated with inhibitors of the PAT, although the signal is weaker in plants treated with L-kynurenin.

**Supplemental Figure 10.**
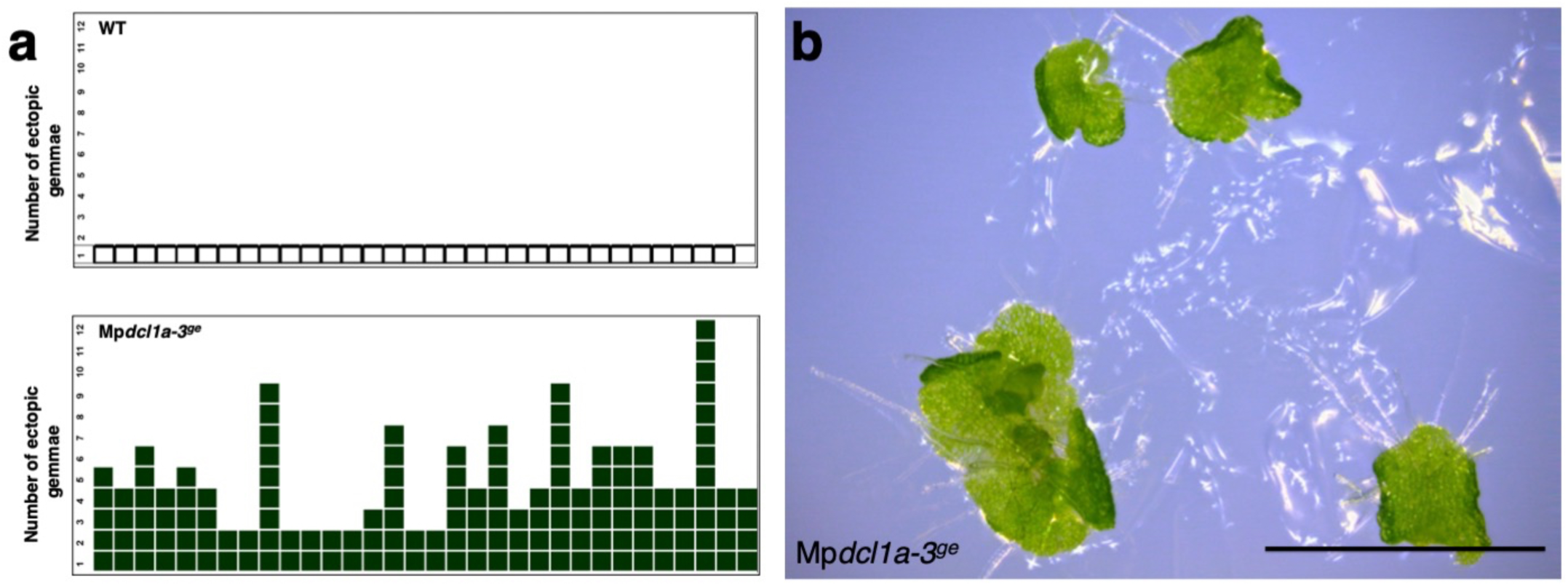
The gemma-on-gemma phenotype of the Mp*dcl1a-3^ge^* mutant is fully penetrant. a) Quantification of the ectopic gemmae phenotype in the Mp*dcl1a-3^ge^* mutant background shows that the phenotype is 100% penetrant. Green colored squares illustrate the presence of ectopic gemmae. (n=32). b) Ectopic gemmae detached from the epidermis of Mp*dcl1a-3^ge^*gemmae and grown in solid media develop into thalli. Scale bar = 1mm.

**Supplementary Figure 11.**
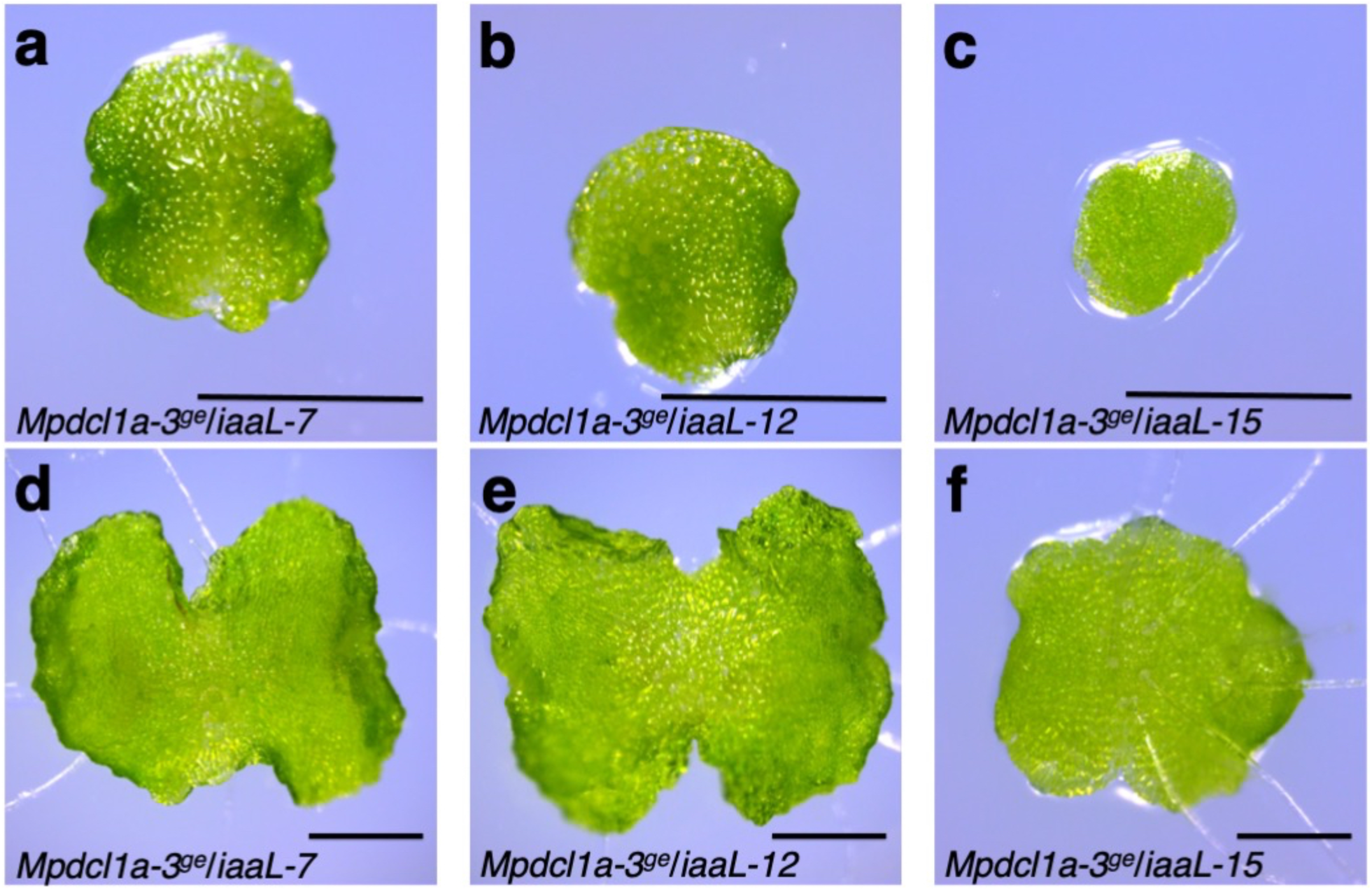
Revertant phenotype of Mp*dcl1a*-*3^ge^*gemmae from plants expressing the auxin conjugating enzyme iaaL. a-c) Revertant phenotype of Mp*dcl1a-3^ge^* gemmae from plants expressing the auxin conjugating enzyme iaaL. d-f) Revertant phenotype of 3-days-old Mp*dcl1a*-*3^ge^* thalli from plants expressing the auxin conjugating enzyme iaaL. Scale bars = 0.5 mm.

**Supplementary Figure 12.**
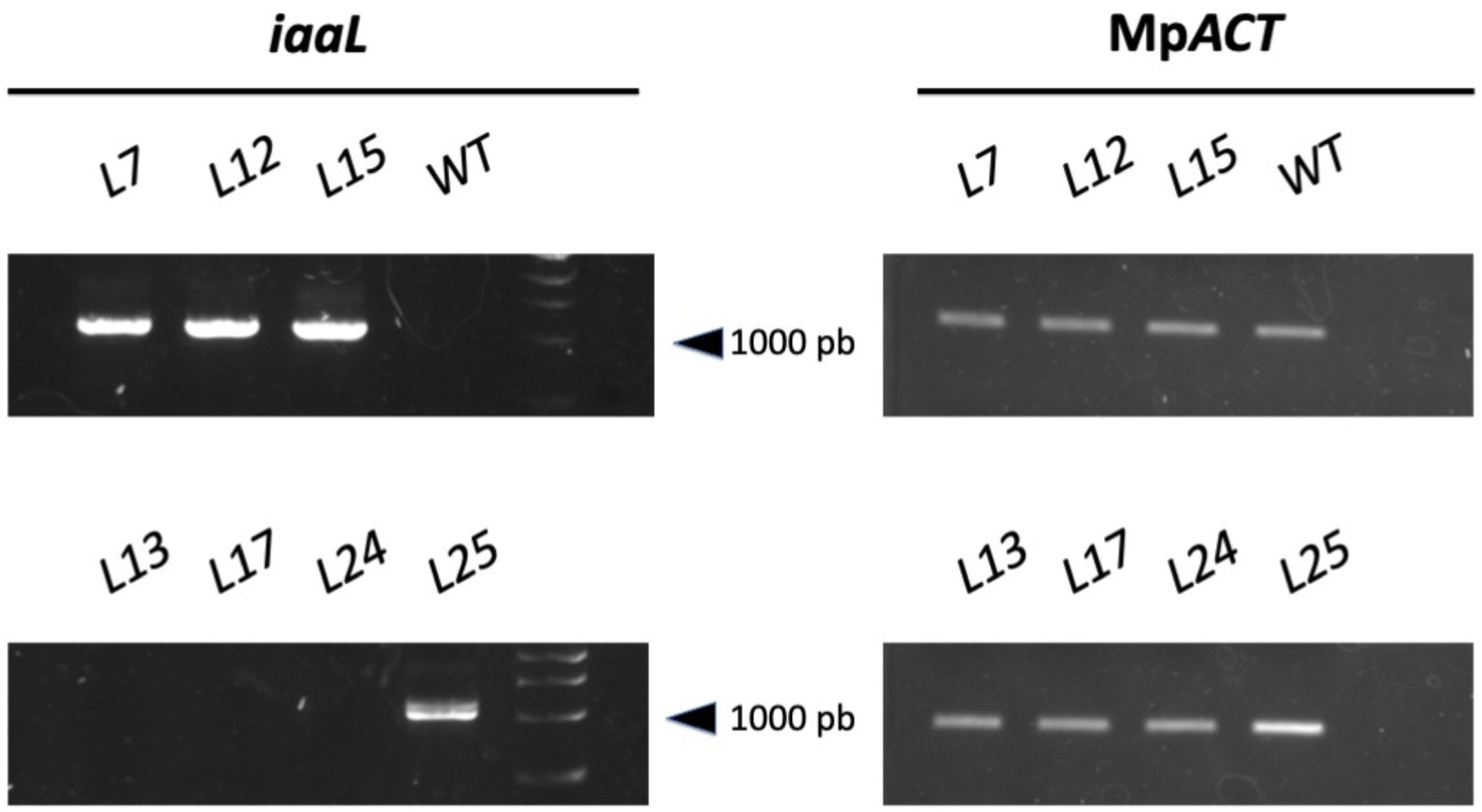
Expression of the auxin conjugator *iaaL* in the absence of the inducer (β-estradiol) in revertant Mp*dcl1a-3^ge^* plants. Semi-quantitative RT-PCR for *iaaL* in uninduced Mp*dcl1a-3^ge^* plants. The upper-left panel shows 1000 bp amplicon for *iaaL* in Mp*dcl1a-3^ge^*revertant lines (L7, L12, L15). The bottom-left panel shows *iaaL* expression (or lack thereof) in 3 of the 4 Mp*dcl1a-3^ge^* lines(L13, L17 and L24) exhibiting ectopic gemmae. The single Mp*dcl1a-3^ge^* line (L25) with ectopic gemmae and positive *iaaL* expression shows lower expression levels than revertant lines. Panels on the right show the expression of Mp*ACT* as a loading control. All experiments were done in the absence of β-estradiol.

**Supplementary Figure 13.**
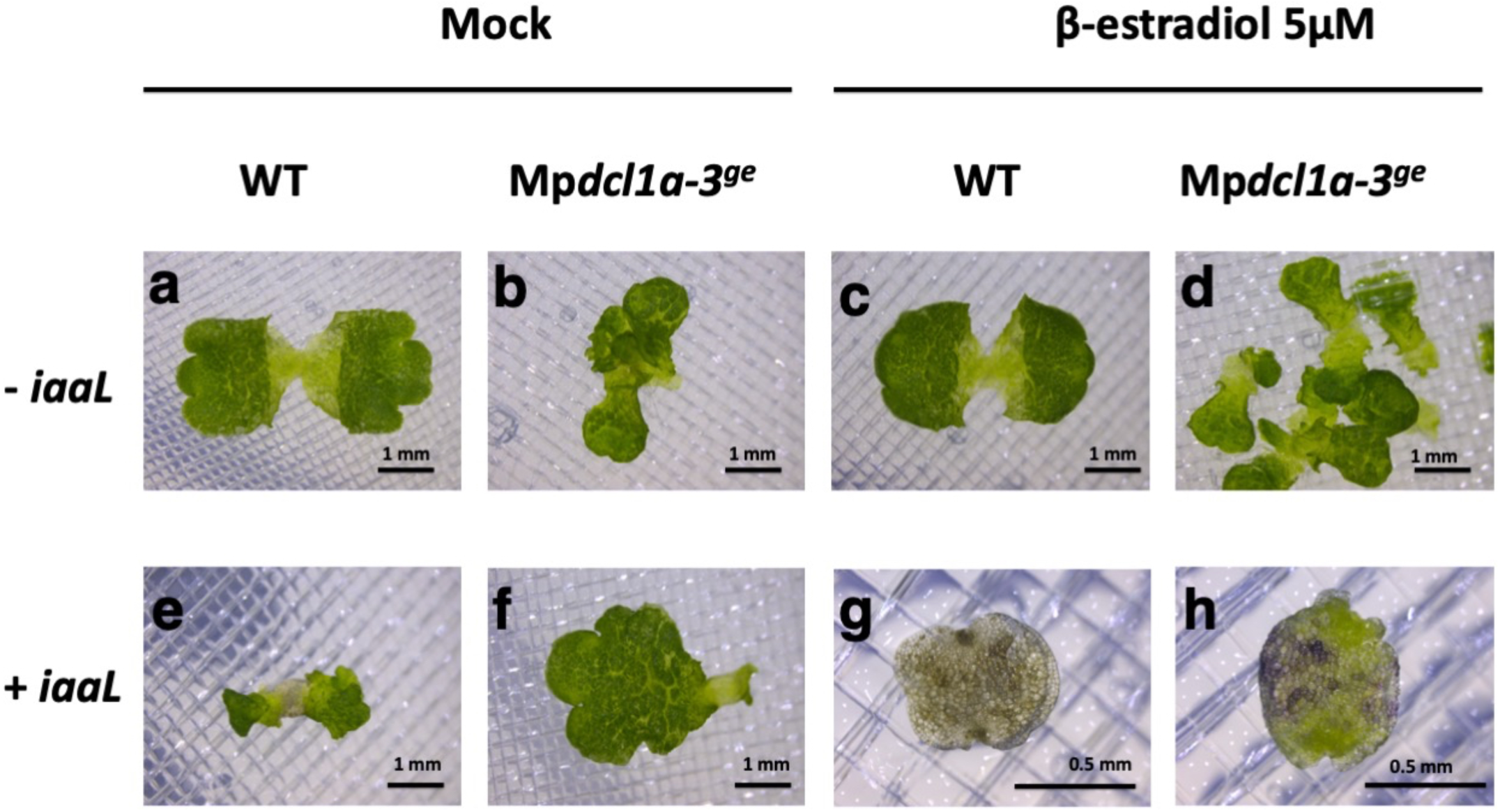
All plants harboring the vector containing the auxin conjugator *iaaL* respond to the inducer β-estradiol. Mock treated or β-estradiol (5μM) treated wild-type (a) and Mp*dcl1a-3^ge^*(b) plants not transformed with the auxin conjugator *iaaL* show no major alterations in their characteristic growth and development. e-f) The majority of mock treated wild-type plants transformed with *iaaL* do not show an evident phenotype (data not shown) but can stochastically exhibit delayed growth and development given the activation of *iaaL* in the absence of β-estradiol since the induction system is leaky. f) The vast majority of mock treated Mp*dcl1a-3^ge^*plants transformed with *iaaL* maintain the mutant phenotype but can stochastically exhibit an improved growth and development in the absence of β-estradiol. β-estradiol treated wild-type and Mp*dcl1a-3^ge^* gemmae display the reported auxin deficient phenotype (Flores-Sandoval, 2015).

